# Chaperone AIP Couples mTORC1 Activation and Catabolic Metabolism During Neonatal Development

**DOI:** 10.1101/2025.09.18.677011

**Authors:** Márta Korbonits, Xian Wang, Sayka Barry, Chung Thong Lim, Oniz Suleyman, Stefano De Tito, Nazia Uddin, Maria Lillina Vignola, Charlotte Hall, Laura Perna, J. Paul Chapple, Gabor Czibik, Sian M Henson, Valle Morales, Katiuscia Bianchi, Viðar Örn Eðvarðsson, Kristján Ari Ragnarsson, Viktoría Eir Kristinsdóttir, Anne Debeer, Yoeri Sleyp, Rena Zinchenko, Glenn Anderson, Michael Duchen, Kritarth Singh, Chih Yao Chung, Yu Yuan, Sandip Patel, Artem O. Borovikov, Hans Tómas Björnsson, Hilde Van Esch, Sharon Tooze, Ezra Aksoy, Caroline Brennan, Oliver Haworth

## Abstract

To grow and divide cells must tightly coordinate anabolic programs with the availability of nutrients and growth factors. This balance is especially critical during postnatal development, when biosynthetic and energetic demands are high, and nutrient supply and neonates have to adapt to periods of fasting. These conditions place acute stress on the proteostasis network, making autophagy essential for nutrient recycling. We found that the chaperone aryl hydrocarbon receptor-interacting protein (AIP) supports both arms of this metabolic balance: promoting anabolic PI3K-AKT signaling for mTORC1 activation and enabling catabolic processes such as proteasomal degradation and autophagy. Loss of AIP causes a severe neonatal metabolic disorder, where affected infants fail to thrive postnatally. Our findings establish AIP as a central regulator of neonatal metabolic adaptation and cellular homeostasis.

**One Sentence Summary:** AIP integrates nutrient sensing and protein recycling to sustain neonatal survival.

## Introduction

For cells to grow and divide, they must rely on anabolic metabolism, a process essential for organismal development. This requires growth factors and an abundant supply of nutrients, which activate the mechanistic target of rapamycin complex 1 (mTORC1), a central metabolic hub that integrates intracellular and extracellular signals^1^. Conversely, during periods of nutrient scarcity, cells engage catabolic processes, including the ubiquitin– proteasome system (UPS) and autophagy-lysosome system (ALS), to recycle macromolecules and sustain anabolic growth^2,3^. The proteostasis network maintains a healthy proteome by coordinating protein synthesis, folding, disaggregation, and degradation^4^. Protein degradation is mediated by the UPS and ALS, with molecular chaperones triaging damaged or misfolded proteins to these pathways^2,3^. During neonatal development, rapid cell proliferation and biomass accumulation impose high biosynthetic and energetic demands on this network^5–7^.

Disruption of proteostasis compromises the cell’s capacity to adapt to dynamic metabolic needs, especially during nutrient deprivation^8–14^. Following birth, neonates undergo periods of fasting and rely on autophagy to maintain a continuous nutrient supply. The inability to induce autophagy during this period is associated with neonatal lethality^10,15^. Despite its importance, the molecular mechanisms that maintain proteostasis during early development remain poorly defined.

Aryl hydrocarbon receptor-interacting protein (AIP) is a ubiquitously expressed, evolutionarily conserved molecular chaperone^16^. Heterozygous germline loss-of-function mutations in *AIP* are linked to pituitary adenomas^17,18^, but its broader metabolic functions are unknown. We present evidence that AIP plays a central role in maintaining both anabolic and catabolic metabolism during periods of high proteostatic demand.

Using an integrative, multi-omics approach combining proteomic, metabolomic, and biochemical analyses of *Aip* knockout mouse embryonic fibroblasts (*Aip* KO MEFs), homozygous AIP-mutant patient-derived dermal fibroblasts (AIPd-PDFs), and AIP-deficient zebrafish, we identify AIP as a critical facilitator of PI3K-AKT driven mTORC1 activation in response to growth factors. Patients born with homozygous *AIP* variants exhibit a severe neonatal disorder, characterized by failure to thrive and postnatal lethality.

Our findings further reveal that AIP sustains proteasome activity, autophagic flux, and lysosomal function during neonatal development, underscoring its central role in coordinating nutrient sensing with cellular growth and protein homeostasis.

### Identification of a new metabolic disease

We have identified five patients from three different countries with a uniform and unique phenotype representing a novel disease: hyperthermia (non-infectious/non-inflammatory), tachycardia, hypercalcemia, severe diarrhea and failure-to-thrive (**Figure 1A-B, Figure S1A**), despite being given state-of-the-art care and feeding support in pediatric intensive care units. The initial two patients, first cousins from a consanguineous family (Family A), died with extreme low body weight and heart failure under the age of 12 months. Whole exome sequencing revealed a biallelic variant of *AIP* in both patients (Family A Member 1 and 2 (FAM1, FAM2)): (c.62G>A); p.(Gly21Asp) (NM_003977.4) (**Figure 1C**, **Table S1**). The identification of further three patients was facilitated by GeneMatcher^19^. Genetic screening revealed a homozygous missense variant in the second family (Family B): c.827C>A; p.(Ala276Glu), while probands from Family C and D, not known to be related but from the same geographical area, had an in-frame deletion of 9 amino acids: c.347_373del; p.Glu116_Val124del (**Figure S1B**). These variants have not been previously described in patients; population frequencies are described in **Table S2**.

**Figure 1.**
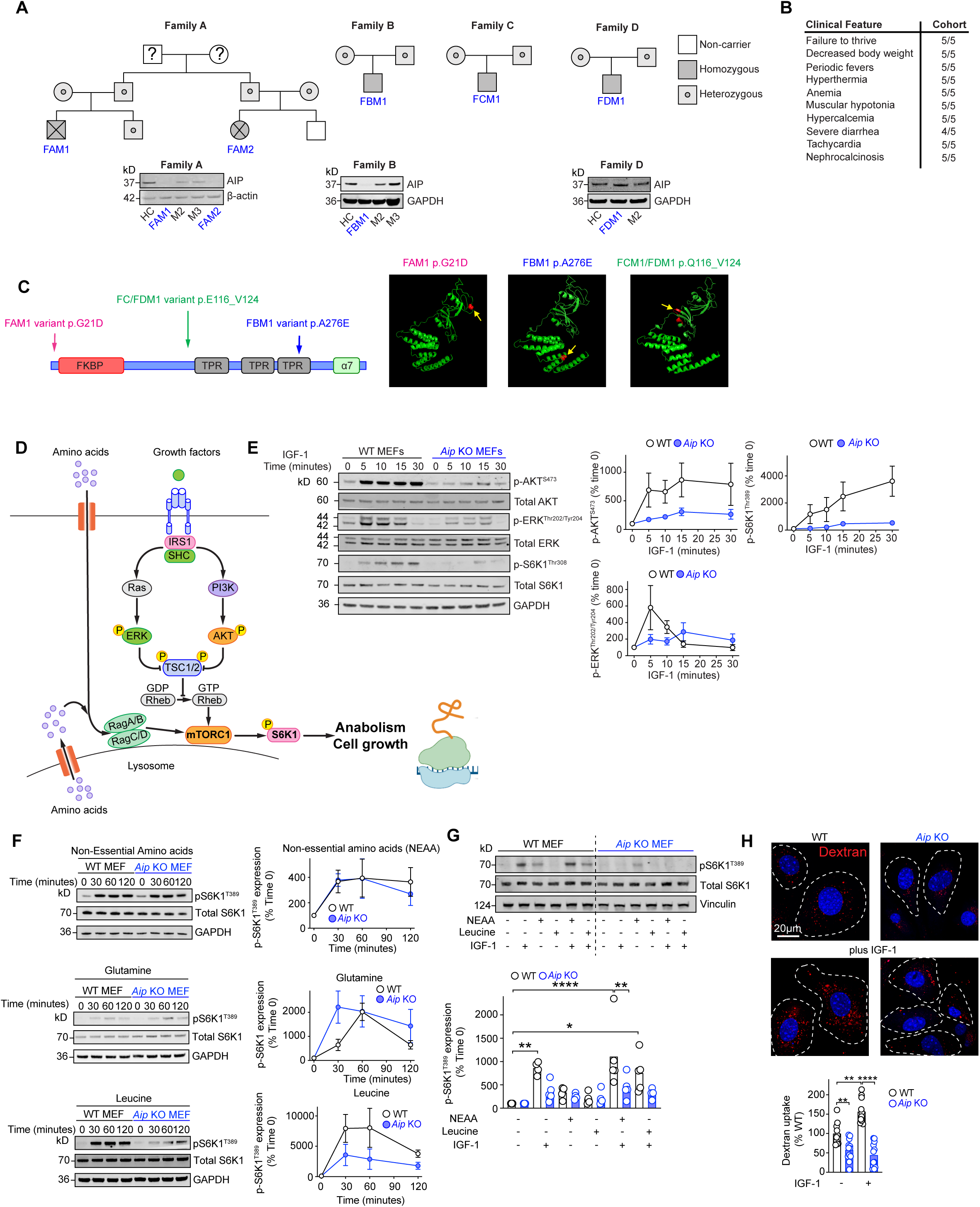
Identification and phenotype of individuals born with AIP deficiency. Family pedigrees of the four families of the children identified to have biallelic AIP variants and Western blot analysis of AIP expression from patient (AIPd-PDFs) and healthy control (HC) derived dermal fibroblasts (A). Salient clinical features found in children born with AIP deficiency (B). Location of the variants in AIP (C). Summary figure of growth factor signaling and mTORC1activation (D). WT and *Aip* KO MEFs were starved overnight in DMEM containing 0.1% serum and stimulated with insulin like growth factor-1 (IGF-1) at 50ng/ml and expression of p-AKT, p-ERK and p-S6K1 examined by Western blotting. (E). WT and *Aip* KO MEFs were starved of amino acids for 2 hours (EBSS with 1% dialyzed FBS) and stimulated with non-essential amino acids (NEAA), leucine (0.8mM) and glutamine (4mM) and mTORC1 (pS6K1^T389^) activation determined by Western blotting (F). WT and *Aip* KO MEFs were starved of amino acids for 2 hours (EBSS with 1% dialyzed FBS) and pre-treated with NEAA, leucine for 15 minutes and then stimulated with IGF-1 and expression levels of p-S6K1 determined (G). Dextran uptake in WT and *Aip* KO MEFs treated with and without IGF-1 (15 minutes) (H). Graphs show the mean ± SEM from at least two independent experiments. 2-way ANOVA with Tukey’s multiple comparison test used for analysis (G-H).

Analysis of AIP protein turnover revealed that two of the three variants, (p.Gly21Asp) and p.(Ala276Glu), resulted in proteins with significantly reduced half-life compared to wild type (WT) AIP (>24 hours), demonstrating that in these patients AIP was rapidly degraded. The in-frame deletion variant (FCM1/FDM1) showed a normal half-life (**Figure S1C**).

### AIP supports PI3K-AKT signaling and mTORC1 activation

To transition from catabolic to anabolic metabolism, cells integrate growth factor and nutrient cues through mTORC1^1^ (**Figure 1D**). Previously, we demonstrated that *Aip* knockout (KO) B cells exhibit impaired AKT activation following IgM stimulation^20^. *Aip* KO MEFs displayed reduced PI3K-AKT and ERK signalling in response to multiple growth factors including insulin-like growth factor 1 (IGF-1), epidermal growth factor (EGF), sphingosine-1-phosphate (S1P), and fetal bovine serum (FBS), resulting in attenuated mTORC1 activation (**Figure 1E** and **Figure S2A-C**). While they responded normally to non-essential amino acids and glutamine, their response to leucine was blunted (**Figure 1F**) and combined leucine/non-essential amino-acids plus IGF-1 stimulation failed to activate mTORC1(**Figure 1G**). In addition, *Aip* KO MEFs showed impaired IGF-1 induced macropinocytosis, limiting nutrient uptake (**Figure 1H**).

The observed defects in growth factor signaling and nutrient uptake in *Aip* KO MEFs led us to hypothesize that these cells increasingly rely on catabolic metabolism to compensate for nutrient insufficiency.

### AIP supports catabolic metabolism

To investigate how AIP deficiency impacts cellular metabolism, we first assessed intracellular amino acid levels using mass spectrometry. Both *Aip* KO MEFs and AIPd-PDFs exhibited significantly reduced levels of amino acids compared to their respective controls (**Figure 2A-B**). To explore the basis of this amino acid deficiency, we performed unbiased Tandem Mass Tag (TMT)-based quantitative proteomics on WT and *Aip* KO MEFs under both nutrient-replete and starved conditions. Principal component analysis (PCA) of the proteomic profiles revealed clear separation between WT and *Aip* KO MEFs, indicating substantial global proteomic changes (**Figure 2C**).

**Figure 2.**
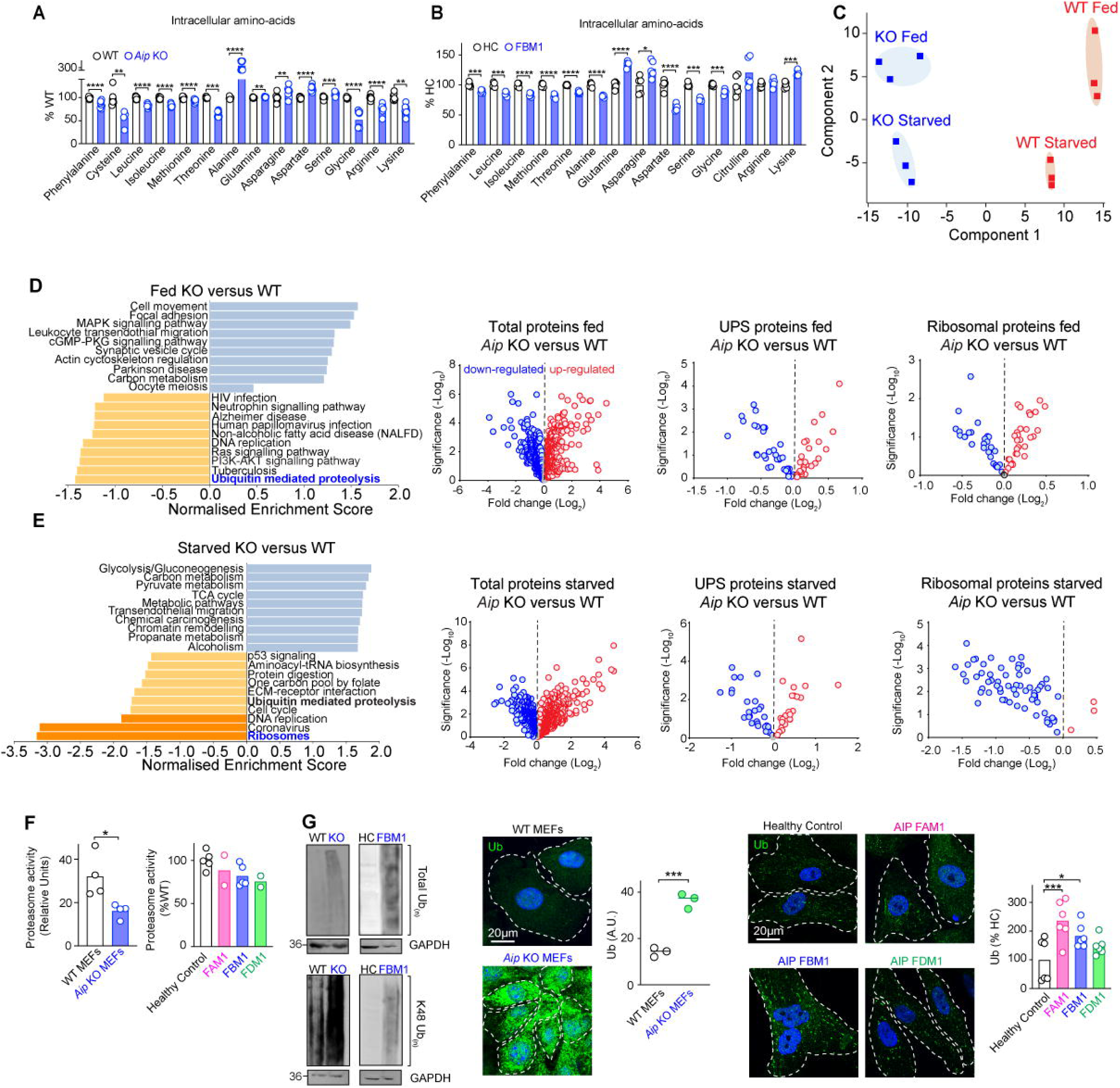
Loss of AIP results in decreased expression of proteins involved in protein recycling. Levels of total amino acids determined by mass-spec analysis from WT and *Aip* KO MEFs (A) and AIPd-PDSFs from HC and FBM1 (B). Proteomic principle component (C). KEGG pathways analysis of TMT-mass spec analysis of WT versus *Aip* KO MEFs under fed and starved conditions showing volcano plots of total proteins, proteins associated with the ubiquitin-proteasome system (UPS)/autophagy-lysosome system (ALS) and ribosomal proteins (D-E). Proteasome activity was measured in WT and *Aip* KO MEFs and HC and AIPd-PDSFs (F). Expression of total ubiquitin and K48 ubiquitin from WT and *Aip* KO MEFs and from HC and AIPd-PDFs examined by Western blotting and by immunofluorescence, arbitrary units (A.U.) (G). Graphs show the mean ± SEM from at least two independent experiments. Student’s un-paired *t*-test (A, B, F), one-way ANOVA (G) used for analysis.

Under fed conditions, the most significantly downregulated process in *Aip* KO MEFs was ubiquitin-mediated proteolysis (**Figure 2D**). Under nutrient-deprived conditions, ribosome biogenesis emerged as the most downregulated process, suggesting that AIP-deficient cells are unable to maintain protein synthesis and biomass accumulation when nutrients are scarce (**Figure 2E**). Given these observations, we next investigated protein recycling mechanisms. Both *Aip* KO MEFs and AIPd-PDFs displayed markedly reduced proteasome activity compared to WT controls (**Figure 2F**). Correspondingly, ubiquitylated proteins accumulated significantly in AIP-deficient cells, indicating a block in protein turnover (**Figure 2G**).

Together, these findings establish AIP as a critical regulator of protein homeostasis and metabolic resilience. Without AIP, cells fail to engage the catabolic programs required to adapt to nutrient stress, leaving them unable to sustain biosynthesis or growth.

### AIP supports the initiation of autophagy

When proteasome function is impaired, autophagy compensates by degrading proteins to maintain proteostasis and energy homeostasis^21–23^. AMPK activation, which occurs during nutrient scarcity, initiates autophagy via phosphorylation of ULK1^24^. Interestingly, *Aip* KO MEFs displayed elevated p-AMPK^T172^ and p-ULK1^S555^ levels even under nutrient-rich conditions, mimicking a starvation-like state (**Figure 3A-B**). This suggests that AIP-deficient cells are metabolically stressed even when nutrients are available, and that the upstream signaling required to initiate autophagy remains functional.

**Figure 3.**
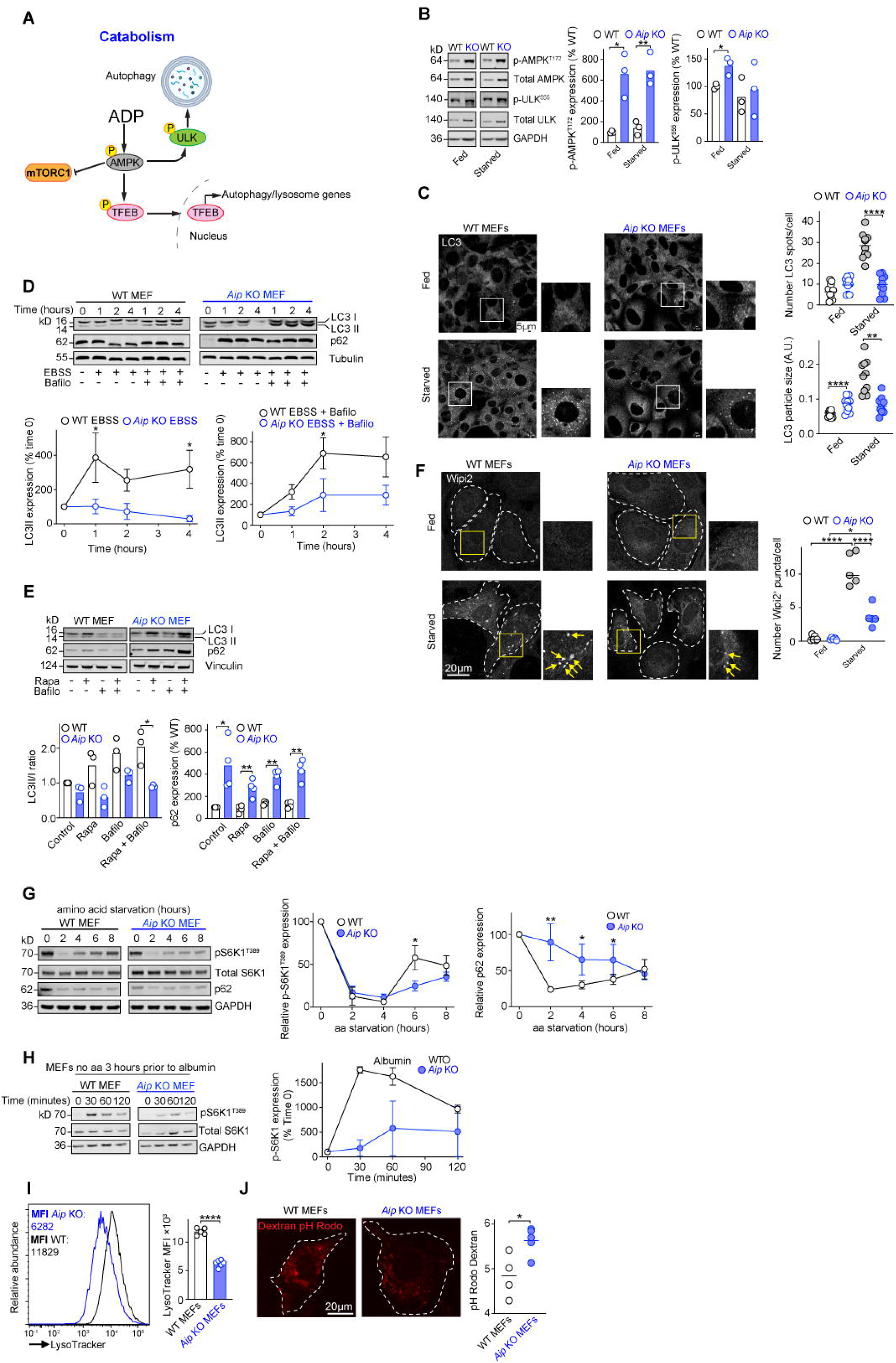
AIP is required to support autophagy. Pathway showing the initiation of starvation-induced autophagy (A). Expression of p-AMPK, p-ULK^555^ in WT and *Aip* KO MEFs cultured under fed and starved conditions (2 hours in Earle’s Balanced Salt Solution (EBSS)) as determined by Western blotting (B). WT and *Aip* KO MEFs cultured under fed and starved conditions and stained for LC3. The size and number of LC3^+^ puncta were determined (C). WT and *Aip* KO MEFs were cultured in EBSS (2 hours) in the presence or absence of bafilomycin A1 (Bafilo) (100nM) for 0, 2 and 4 hours and the expression of LC3I/II and autophagic flux determined (D). WT and *Aip* KO MEFs were treated with Rapamycin (Rapa) (100 nM) and Bafilo (100nM) for 2 hours and the expression of LC3I/II and p62 determined (E). Wipi2^+^ puncta in WT and *Aip* KO MEFs under fed and starved (2 hours EBSS) conditions (F). WT and *Aip* KO MEFs were starved of amino acids (EBSS media supplemented with 1% dialyzed FBS) and mTORC1 (pS6K1^T389^) reactivation over time and p62 expression determined by Western blotting (G). Expression of lysotracker in WT and *Aip* KO MEFs determined by flow cytometry (I). pH of endosomes determined by dextran-pH Rodo in WT and *Aip* KO MEFs (J). WT and *Aip* KO MEFs were starved of amino acids for 2 hours and given albumin and mTORC1 (pS6K1^T389^) expression determined by Western blotting (H). Expression of lysotracker in WT and *Aip* KO MEFs determined by flow cytometry (I). pH of endosomes determined by dextran-pH Rodo in WT and *Aip* KO MEFs (J). Graphs show the mean ± SEM from at least two independent experiments. Student’s un-paired and paired *t*-test (B, C, E, F, I, J) 2-way ANOVA with Tukey’s multiple comparison test used for analysis (D, G)

However, starvation failed to increase LC3-II levels in *Aip* KO MEFs (**Figure 3C**). Inhibition of autophagosome–lysosome fusion with bafilomycin A1 did not further increase LC3-II accumulation, indicating that the block in autophagy occurs upstream of autophagosome fusion (**Figure 3D**). Similarly, rapamycin, an mTORC1 inhibitor and autophagy inducer, failed to promote autophagy or lead to a reduction in p62 expression in *Aip* KO MEFs (**Figure 3E**). These results point to a defect in autophagy initiation. Functionally, *Aip* KO MEFs displayed reduced viability during prolonged nutrient deprivation compared to wild-type controls, and viability was further decreased by bafilomycin treatment, reflecting impaired catabolic capacity (**Figure S3A**). Importantly, lentiviral re-expression of human AIP in *Aip* KO MEFs restored LC3-II induction and reduced ubiquitin accumulation, confirming the specificity of the defect (**Figure S3B**).

Autophagy initiation depends on PI3P and PI4P signalling at omegasomes^25^. While *Aip* KO MEFs maintained near-normal PI3P levels (**Figure S3C**), they failed to recruit Wipi2 upon starvation (**Figure 3F**), indicating a block in downstream ATG protein assembly^26,27^. PI4P dynamics were also impaired, *Aip* KO MEFs had fewer PI4P+ and PI4P+/Wipi2+ puncta (**Figure S3D**), and PI4K3β failed to re-localise from the perinuclear region to autophagy initiation sites under starvation (**Figure S3E**). These defects implicate AIP in coordinating PI4P-dependent trafficking of autophagy regulators.

At the transcriptional level, *Aip* KO MEFs displayed constitutive activation of TFEB^28^ and TFE3 (**Figure S4A-B**), driving elevated basal expression of autophagy genes such as *Atg9b*, *Ctsl1*, and *Tfeb* (**Figure S4C**). However, this compensatory program could not restore autophagic induction, as both MEFs and AIP-deficient fibroblasts (AIPd-PDFs) failed to further upregulate autophagy upon starvation (**Figure S5A-D**).

In summary, AIP-deficient cells display chronic activation of starvation signalling yet fail to initiate effective autophagy, owing to defective PI4P production, impaired PI4K3β trafficking, and blocked recruitment of initiation machinery. Thus, AIP emerges as a key regulator of early autophagy, indispensable for cellular adaptation to nutrient stress.

### AIP is required for healthy lysosome function

Lysosomes serve as metabolic hubs, coordinating nutrient sensing, macromolecular degradation, and mTORC1 signalling^29^ ^30^. EM analysis of *Aip* KO MEFs and AIPd-PDFs revealed increased autophagosomes, reduced lysosomes, and abnormal vesicular structures, consistent with defective lysosomal function and trafficking (**Figure S6A-B**). Autophagy turnover was impaired, as evidenced by delayed mTORC1 reactivation and p62 degradation following starvation, suggesting defective autophagic lysosome reformation^26,31^ (**Figure 3G**). Refeeding with albumin failed to promptly restore mTORC1 activity, further indicating compromised lysosomal proteolysis (**Figure 3H**). Mechanistically, *Aip* KO MEFs displayed reduced LysoTracker staining (**Figure 3I**) and elevated lysosomal pH (∼5.6) compared to WT MEFs (∼4.8), confirming impaired acidification and loss of degradative capacity (**Figure 3J**).

Transcriptomic profiling of *Aip* KO MEFs revealed downregulation of multiple lysosome-related genes, including those associated with lysosomal storage diseases (LSDs)^29^ (**Figure S7A-B**). Proteomic analysis further supported lysosomal dysfunction, showing significant reduction in key lysosomal enzymes such as acid alpha-glucosidase (GAA) and Niemann-Pick disease type C2 (NPC2) (**Figure S7C**). These enzymes are essential for glycogen and lipid degradation^29^, respectively and their deficiency provides evidence that AIP-deficient lysosomes are functionally impaired, resulting in ineffective protein catabolism and blunted mTORC1 reactivation.

Together, these results demonstrate that AIP is required for the formation of functionally competent lysosomes, enabling proper acidification, cargo degradation, and nutrient recycling.

### AIP Is Required to Support Neonatal Metabolic Reprogramming

Neonatal life is marked by high energy demands and rapid growth, requiring a fundamental shift in metabolic programs to adapt to the postnatal nutrient environment^7^. Although *Aip* KO MEFs appeared outwardly normal and proliferated under standard culture conditions, we hypothesized that these cells employ alternative metabolic strategies to compensate for impaired protein recycling and sustain anabolic growth.

Metabolomic profiling revealed a strong reliance on glycolysis in *Aip* KO MEFs, as evidenced by elevated levels of glycolytic end-products including pyruvate, lactate, and alanine (**Figure S8A-F**). Tracer studies with U-¹³C₆ glucose revealed increased incorporation of glucose-derived carbon (m+3) into glucogenic amino acids including alanine, serine, aspartate, asparagine, and glutamate, demonstrating a preferential diversion of glucose from oxidative TCA metabolism toward amino acid biosynthesis, thereby shifting metabolism from energy production to cataplerosis (**Figure 4A-B**).

**Figure 4.**
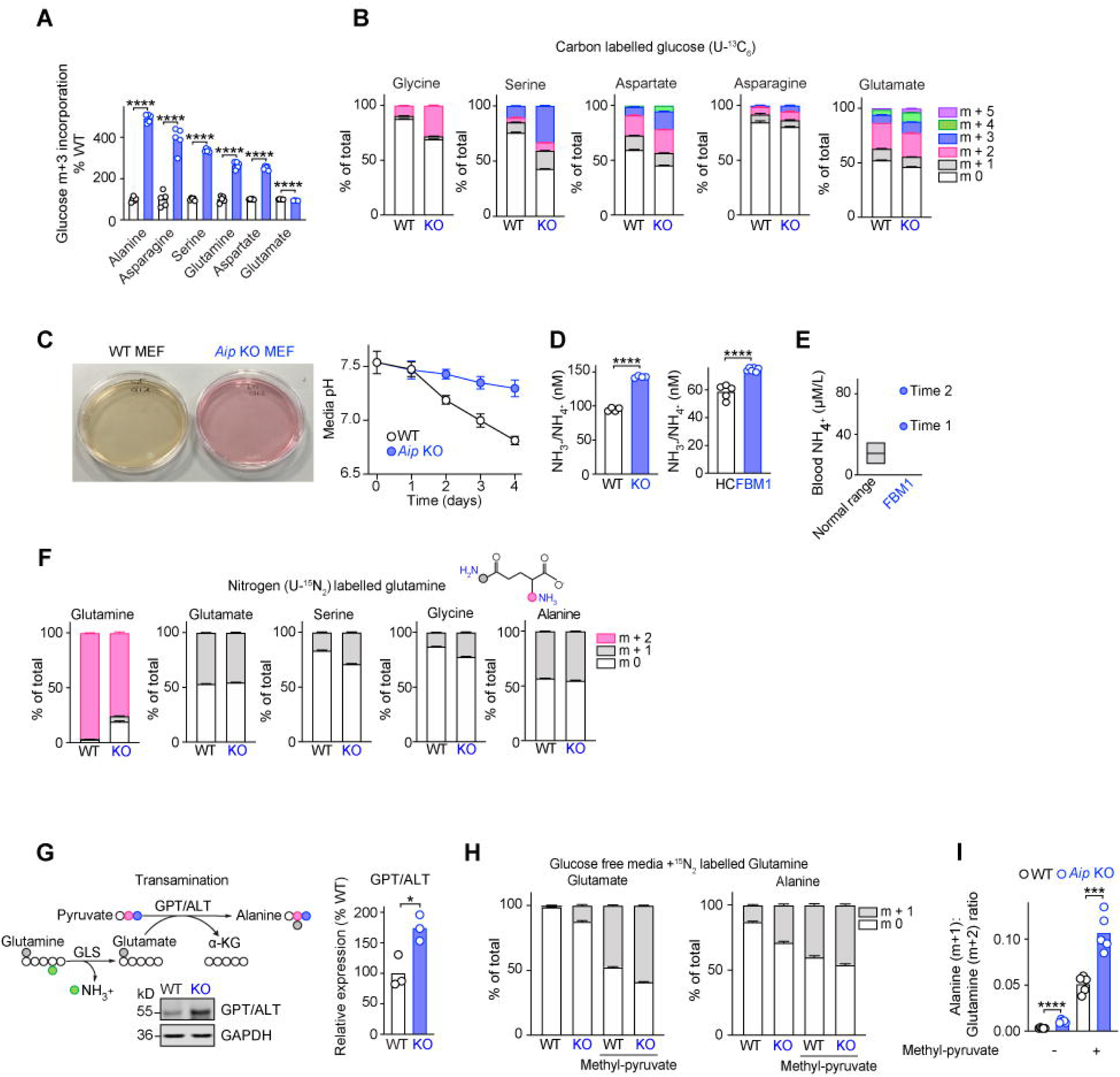
AIP-deficient cells undergo metabolic reprogramming. Relative amount of m+3 universally labelled glucose (U-^13^C_6_) incorporated into amino acids from WT and *Aip* KO MEFs (A). Relative proportion of uniformly labelled glucose in amino acids from WT and *Aip* KO MEFs (B). Media pH over time in WT and *Aip* KO MEFs (C). Levels of NH_3_^+^/NH_4_ ^+^ in WT and *Aip* KO MEFs and HC and AIPd-PDFs (D). Elevated blood NH_4_ levels determined from patient FBM1 compared to the normal range (E). Relative levels of glutamine metabolites in WT and *Aip* KO MEFs and relative levels of glutamine metabolites and flux derived from uniformly labelled (U-^13^C_6_) glucose in WT and *Aip* KO MEFs (F). Transamination of glutamine using pyruvate as an amino acceptor by the enzyme GPT/ALT expressed in WT and *Aip* KO MEFs (G). WT and *Aip* KO MEFs were cultured in glucose free media with U-^15^N_2_ glutamine in the presence or absence of 10mM methyl-pyruvate and the relative proportion of U-^15^N_2_ glutamine isotopologues determined (H). The ratio of alanine (m+1): glutamine (m+2) (I). Graphs show the mean ± SEM from at least two independent experiments. Student’s un-paired *t*-test used for analysis (A, D, G, I).

Consistent with this glycolytic bias, both *Aip* KO MEFs and AIPd-PDFs displayed hypersensitivity to glucose deprivation (via 2-deoxy-D-glucose) and ATP synthase inhibition (via oligomycin) (**Figure S8G-H**), underscoring a compensatory reliance on glycolysis for both energy and precursor supply. Moreover, *Aip* KO MEFs showed greater incorporation of glucose-derived carbon (m+3) into TCA cycle intermediates, including aconitate, α-ketoglutarate (α-KG), and malate (**Figure S9A-C**), suggesting increased carbon flux into the TCA cycle despite compromised oxidative phosphorylation. Interestingly, *Aip* KO MEFs exhibited increased mitochondrial membrane potential and volume (**Figure S9D-F**), elevated mitochondrial NADH levels, and reduced ATP production (**Figure S9G-H**), pointing to mitochondrial dysfunction characterized by inefficient oxidative phosphorylation despite expanded mitochondrial biomass.

Given the accumulation of ubiquitinated proteins in AIP-deficient cells (**Figure 2G**), we hypothesized that increased amino acid deamination may serve dual roles, detoxifying excess protein cargo and supplying carbon skeletons for anaplerosis. Supporting this, unlike WT MEFs, *Aip* KO MEFs failed to acidify their culture medium (**Figure 4C**), suggesting altered ammonia metabolism. Indeed, *Aip* KO MEFs and AIPd-PDFs accumulated significantly higher levels of ammonia (NH₃⁺/NH₄⁺) (**Figure 4D**). These findings, combined with reduced intracellular and extracellular amino acid levels (**Figure 2A-B**, **Figure S8E**), indicate enhanced amino acid catabolism with inadequate nitrogen clearance. This hypothesis was confirmed *in vivo* by elevated blood ammonia in patient FBM1 on two clinical occasions (**Figure 4E**), highlighting a conserved role for AIP in nitrogen metabolism. Further, U-¹³C₅ glutamine tracing revealed impaired conversion to α-KG, indicating defective oxidative and reductive glutaminolysis (**Figure S10A**). Labelling with U-¹LN₂ glutamine showed increased nitrogen transfer into serine, glycine, and alanine (**Figure 4F, Figure S10B**), suggesting nitrogen detoxification via transamination.

Our data indicates that, in the absence of effective nitrogen recycling, transaminated amino acids serve as alternative nitrogen sinks, thereby reducing the availability of carbon backbones to be utilized for anaplerosis and energy production. Consistently, glutamic-pyruvate transaminase (GPT)/alanine aminotransferase (ALT), an enzyme that transfers amino groups from glutamine to pyruvate to form alanine, was upregulated in *Aip* KO MEFs (**Figure 4G**). In glucose-free media supplemented with U-¹LN₂ glutamine, *Aip* KO MEFs incorporated significantly more nitrogen into alanine (m+1) than WT MEFs, particularly with methyl-pyruvate supplementation (**Figure 4H-I**). This alanine shunt was also reflected in elevated serum alanine levels in patients FAM1 and FBM1 (**Figure S10C**), reinforcing its physiological relevance.

Collectively, these findings show that AIP deficiency disrupts neonatal metabolic reprogramming by impairing energy production, increasing dependence on glycolysis and transamination, leading to toxic nitrogenous waste accumulation. While these adaptations may transiently support cell survival, they do so at the cost of anabolic growth, positioning AIP as a critical regulator of neonatal metabolic homeostasis.

### AIP loss in zebrafish recapitulates patient and MEF phenotype

To assess the in vivo relevance of AIP in vertebrate development and metabolism, we generated *aip* KO zebrafish using CRISPR-Cas9 gene editing (**Figure S11A-B**). In agreement with the phenotype observed in children with biallelic AIP deficiency, homozygous *aip* KO zebrafish exhibited reduced growth and failure to thrive, with significantly smaller body size by 6 days post-fertilization (dpf) (**Figure 5A**). Notably, survival between WT and *aip* KO fish was comparable up to 6 dpf, but *aip* KO fish exhibited a dramatic drop in survival, coinciding with depletion of the yolk, a critical nutrient source during early development (**Figure 5B-C**).

**Figure 5.**
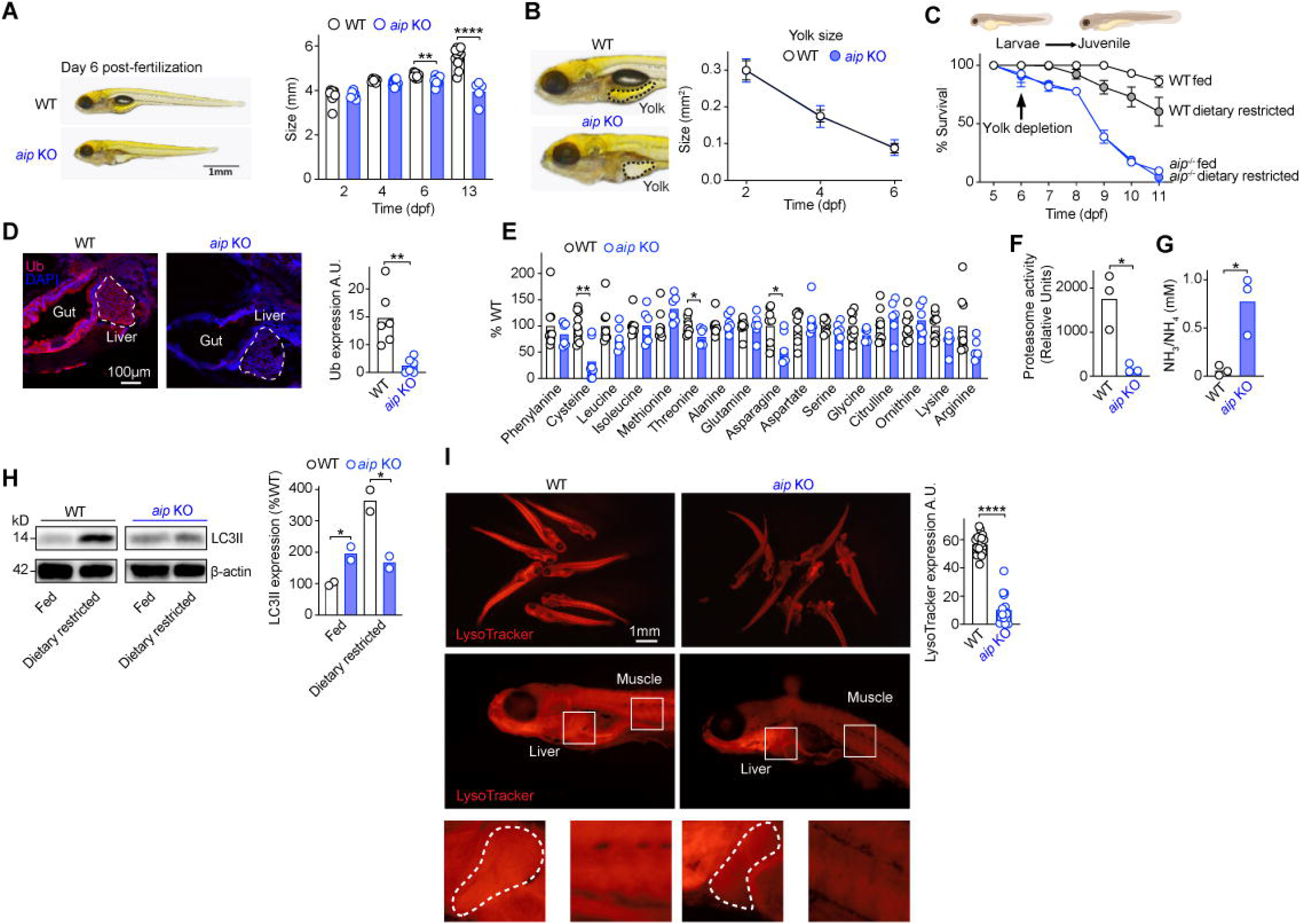
*aip* knockout zebrafish recapitulate the human and phenotype. Size of WT and *aip* knock out zebrafish (KO) from day 2 dpf (A). Comparison of yolk size from day 2 to day 6 post fertilization (dpf) in WT and *aip* KO zebrafish (B). Survival rate of WT and *aip* KO zebrafish from day 5 post fertilization (dpf) under fed and dietary restricted conditions. (C). Ubiquitin expression examined by immunofluorescence (D). Relative amino acids from WT and *aip* KO zebrafish (E). Relative proteasome activity in WT and *aip* KO zebrafish (F). Ammonia (NH_3_^+^/NH_4_^+^) concentration from WT and *aip* KO zebrafish (G). LC3 expression under fed and dietary restricted conditions (H). LysoTracker expression in WT and *aip* KO zebrafish (I). Fish used for experiments were between 5-13 dpf. Graphs show the mean ± SEM from at least three independent experiments. 2-way ANOVA with Tukey’s post hoc test analysis (A), Student’s un-paired *t*-test used for analysis (D-I).

In contrast to *Aip* KO MEFs, which accumulate ubiquitylated proteins under nutrient-rich *in vitro* conditions, *aip* KO zebrafish exhibited reduced levels of ubiquitinated proteins, likely reflecting insufficient nutrient uptake *in vivo* and a physiological downscaling of translation in an attempt to economize (**Figure 5D**). Biochemical analysis of *aip* KO fish revealed decreased levels of several amino acids (**Figure 5E**), reduced proteasome activity (**Figure 5F**), and elevated ammonia levels (**Figure 5G**), mirroring the findings in *Aip* KO MEFs and AIPd-PDFs. These metabolic changes suggest impaired proteostasis and nitrogen clearance *in vivo*.

Given that autophagy is essential for the larval-to-juvenile transition in zebrafish and for neonatal survival in mammals^10–13^, we next assessed autophagic capacity in *aip* KO larvae. Upon dietary restriction, *aip* KO zebrafish failed to upregulate LC3, and baseline LC3 expression was markedly reduced (**Figure 5H**). In line with observations in *Aip* KO MEFs, lysosomal function was compromised, as shown by reduced LysoTracker staining (**Figure 5I**), further confirming lysosomal dysfunction as a hallmark of AIP deficiency.

To investigate metabolic reprogramming *in vivo*, we treated 4 dpf zebrafish larvae with U-¹³C₆-glucose or U-¹³C₅-glutamine. Although the magnitude of labelling was generally lower than in cultured MEFs, *aip* KO fish displayed elevated levels of pyruvate and its aminated/reduced forms, along with a greater contribution of glucose-derived carbon to TCA intermediates and amino acid synthesis, being used for anaplerosis rather than full catabolism towards ATP and amino acid synthesis, consistent with ammonia scavenging (**Figure S12A-C**).

In summary, AIP-deficient zebrafish faithfully recapitulate key aspects of the metabolic, proteostatic, autophagic and lysosome defects observed in patient cells and *Aip* KO MEFs. These findings establish zebrafish as a robust *in vivo* model for studying the metabolic consequences of AIP loss and underscore the essential role of AIP in supporting early post-embryonic development and nutrient adaptation.

## Discussion

Protein synthesis is an energetically demanding process, particularly in rapidly growing cells, and requires a constant supply of amino acids and cellular energy^6^. This process is tightly regulated by mTORC1, which integrates inputs from nutrients, growth factors, and cellular energy status to promote anabolic metabolism. mTORC1 activation occurs only under conditions of sufficient nutrient and growth factor availability and is essential for normal growth and development^14^. During the transition from fetal to neonatal life, cells experience a metabolic shift that demands coordination between anabolic and catabolic processes to ensure sustained nutrient and amino acid availability. Our data reveal that the chaperone protein AIP is essential during this critical developmental window, coordinating the balance of catabolic and anabolic processes allowing cellular and organismal growth, acting to support proteostasis by facilitating proteasome activity and ensuring efficient function of the autophagy-lysosome system.

For cells to commit to growth and cell division, mTORC1 needs to be activated by both growth factors and the availability of sufficient nutrients^14^. Mechanistically, *Aip* KO MEFs were unresponsive to stimulation by multiple extracellular cues including IGF-1, EGF, S1P, and serum via both growth factor receptors and G-protein-coupled receptors. This unresponsiveness affected both PI3K-AKT and RAS-ERK signaling pathways, suggesting that AIP plays a critical role in supporting early, proximal signal transduction across multiple receptor pathways. In this context, *Aip* KO MEFs reduced activation of the PI3K-AKT pathway in response to growth factors, leading to impaired nutrient uptake and a compensatory metabolic shift toward catabolic processes such as autophagy to maintain cellular energy balance.

AIP-deficient cells are more reliant on cataplerosis as a consequence of AIP-deficient cells failing to adapt to nutrient deprivation, owing to compromised proteasome and lysosomal function. During postnatal metabolic stress, when nutrient demand is high, failure to recycle proteins through the UPS and ALS severely compromises cellular function. These defects hinder mTORC1 activation, impair protein biosynthesis, and prevent biomass accumulation, contributing to growth failure. With both major proteolytic systems compromised, AIP-deficient cells adopt alternative metabolic strategies to manage amino acid scarcity while detoxifying accumulated proteins. Notably, ribosome biogenesis, an energetically expensive process which consumes approximately 50% of a cell’s total transcriptional and translational capacity^32^, is drastically reduced in *Aip* KO MEFs during starvation, reflecting a broader downscaling of anabolic processes and translational output.

Clinically, biallelic AIP deficiency in children leads to a severe metabolic disease that is unresponsive to conventional interventions. These children are born with normal body size parameters but exhibit postnatal growth failure, suggesting that AIP is dispensable for *in utero* development but essential for neonatal adaptation. This is further supported by our *aip* KO zebrafish model, which showed normal survival until yolk depletion, after which the fish failed to survive the transition to exogenous feeding and post-larval development. These parallels highlight the evolutionary conservation of AIP’s function in supporting neonatal survival and metabolic adaptation.

Postnatal life demands a rapid metabolic reprogramming, from passive nutrient delivery from the maternal blood supply to active nutrient uptake and recycling. Organisms must efficiently balance catabolic and anabolic metabolism to cope with intermittent nutrient availability and support high biosynthetic demands. The striking phenotype observed across AIP-deficient cells, zebrafish, and human patients underscores AIP’s crucial role during this transition. AIP serves as a key integrator of proteostasis and metabolism, enabling growth by ensuring both sufficient nutrient mobilization and biosynthetic capacity (**Figure S13**).

Together, our findings position AIP as an unrecognized essential regulator of neonatal metabolic reprogramming, acting as a critical hub of nutrient sensing, protein recycling, and growth signaling. These insights may inform future therapeutic approaches for metabolic diseases associated with proteostasis dysfunction.

## Methods

### Identification of homozygous variants in AIP

Whole exome sequencing was performed for Family A using SeqCap EZ Exome Probes v3.0 (Roche) on an Illumina HiSeq2500 system with an average depth of coverage of 125×. Given the consanguineous pedigree and the similar phenotype in both probands, an autosomal recessive condition was suspected. Genetic analysis focused on homozygous variants in exonic regions or splice sites with an ultra-rare allele frequency in the population (allele frequency <0.001%) (using gnomAD version 4.1.0) that are shared between the probands. Whole genome sequencing analysis was performed in Family B due to the lack of diagnosis for the severe abnormalities of the proband using DeCode genetics (https://www.decode.com). Whole exome sequencing was performed for Family C and D probands using the Illumina TruSeq® ExomeKit (Illumina, San Diego, CA, United States) and custom oligonucleotides (IDT xGen Exome Research Panel v1.0) included coding regions of over 20,000 protein-coding genes. Paired-end sequencing (2 × 150 bp) was carried out on an Illumina NextSeq 500. The sequencing data were processed using Illumina’s Basespace software (Enrichment 3.1.0). Variant filtering was based on their frequency, with variants having a frequency of less than 1% in The Genome Aggregation Database (gnomAD v.2.1.1). In all cases, Sanger sequencing was used to confirm the variants in probands and family members.

### Cell culture

MEFs (ATCC) and human dermal fibroblasts (from biopsy of the lower arm) were grown in high glucose (25mM) DMEM supplemented with 10% FBS and penicillin and streptomycin (Thermo Fisher Scientific) unless otherwise stated for experimental purposes. Cells were regularly checked to ensure that they were mycoplasma negative. To induce autophagy, cells were washed 3× in PBS and Earle’s Balanced Salt Solution (EBSS) was added to cells for 2 hours unless otherwise stated.

### Protein half-life analysis

*In vitro* protein half-life analysis was performed as previously described ^33^. *AIP* variants were created using site-directed mutagenesis on MYC-tagged pcDNA3.0 plasmid vector containing the human *AIP* sequence (NM_003977.4). Primers are available on request. The wild-type and variant AIP vectors were transfected into HEK293 cells using lipofectamine 3000 (Thermo Fisher Scientific). 24 hours later, transfected HEK cells were treated with 50µg/ml cycloheximide (CHX) for 0, 6, 12 and 24 hours and Western blot analysis performed.

### Cell viability

Cells (40,000/well) were plated in triplicate in a 96-well plate. The viability of cells the day after plating (time 0) and indicated intervals thereafter was measured using a cell viability kit (CellTiter-Glo, Promega). Viability was calculated as a percentage compared to day 0.

### qPCR

RNA was extracted from cells following treatment using TRIzol (Invitrogen). cDNA was made using High-Capacity cDNA reverse transcription kit (Applied Biosystems) and used with Brilliant III UltraFast SYBR green qPCR master mix (Agilent) on the Agilent AriaMx PCR system. The PCR cycling program implemented was 1 cycle of denaturation for 3 minutes at 95°C, proceeded by 40 cycles of denaturation for 20 seconds at 95°C, and annealing for 20 seconds at 60°C.

### Dextran uptake

Cells (50,000) were added to coverslips and the next day, media was removed, and cells were washed three times in PBS and incubated overnight in complete media containing 0.1% FBS. Cells were stimulated with IGF-1 (50ng/ml) for 15 minutes before the addition of dextran-Texas red (Thermo Fisher Scientific) used at 1mg/ml and incubated for 15 minutes. Cells were washed three times in PBS, followed by the addition of complete media with 10% FBS and pulsed for 30 minutes. Cells were fixed and examined by confocal microscopy and data analyzed using Image-J Software with the mean of 3-6 separate images (30-60 cells) used with the same threshold applied across all samples.

### Measuring MitoTracker expression

MEFs (50,000/well) were seeded onto a 24-well plate on the day prior to the experiment. On the day of the experiment, MEFs were washed 2× in PBS before being incubated with Mitosox/Mitotracker Green/Mitotracker Red according to the manufacturer’s instructions. Samples were acquired on an LSR Fortessa (BD Biosciences) and the data analyzed using FlowJo software version 9.3.1 (Tree Star, Inc, Ashland, USA).

### Measuring glucose uptake using 2-NBDG

MEFs (50,000/well) were seeded onto a 24-well plate on the day prior to the experiment. On the day of the experiment, MEFs were washed 2× in PBS before being incubated with glucose-free DMEM supplemented with 10% dialyzed FBS for 1 hour at 37°C. Media was removed and DMEM with 100µM 2-NBDG (2-(*N*-(7-Nitrobenz-2-oxa-1,3-diazol-4-yl)Amino)-2-Deoxyglucose) added to MEFs and incubated for 30 minutes at 37°C. Media was then removed, and the MEFs were washed with PBS before adding trypsin to harvest the cells in preparation for analysis of 2-NBDG uptake by flow cytometry.

### Measuring NAD/NADH ratio

This was performed as described previously ^34^.

### Measuring mitochondrial membrane potential

MEFs (10,000/well) were seeded onto glass-bottom 24-well plates, 2-3 days before imaging. Media was replaced with phenol red free DMEM supplemented with 1mM glucose, 1mM glutamine, 10mM HEPES, adjusted to pH 7.4 and incubated with 25nM tetramethylrhodamine methyl ester (TMRE) for 30 minutes at 37 °C. Cells were imaged with an LSM 880 (Carl Zeiss) confocal microscope using Fluorescent 63×/1.40 oil immersion objective lens at 37°C. TMRE was excited with a 561nm Argon laser with an output power of 0.2mW as described in ^34^.

### Measuring pH using pHrodo

MEFs (100,000/well) were seeded onto 35mm MatTek dishes the day prior to the experiment. pH calibration was established using pHrodo Red AM prepared in cell loading solution (Thermo Fisher Scientific) supplemented with 20µM nigericin and 20µM valinomycin adjusted to different pH values (4.5, 5.5, 6.5, 7.5). On the day of the experiment, MEFs were washed 2× in live cell imaging solution (Thermo Fisher Scientific) and incubated with the pH adjusted pHrodo solutions for 5 minutes at 37°C before live cell imaging to produce a calibration curve. Cells were incubated with dextran pHrodo for 15 minutes at 37°C before live cell imaging.

### DQ-OVA/LysoTracker

MEFs (50,000/well) were seeded onto coverslips the day prior to the experiment. DQ-OVA (Thermo Fisher Scientific) was prepared according to the manufacturer’s instructions and incubated with cells for 15 minutes at 37°C before removing and washing the cells 2× in PBS and adding fresh media for 30 minutes before adding LysoTracker for 15 minutes.

### Amino acid, glutamine, glucose withdrawal

MEFs were washed 3× in PBS and media without amino acids (EBSS) added. For amino acid stimulation, cells were starved of amino acids for 3 hours and glutamine (4mM final concentration), leucine (0.8mM final concentration) added in the presence of 3% dialyzed FBS for the indicated times.

### Metabolomic analysis

MEFs (10^6^/well) were seeded onto 6-well plates (5× replicates and 1 for cell counting) and the next day MEFs were washed 3× in glucose-free media and 2mL glucose-free media supplemented with 10% dialyzed FBS with 25mM uniformly labelled glucose (U-^13^C_6_) or 4mM uniformly carbon labelled glutamine (U-^13^C_5_) or nitrogen labelled glutamine (U-^15^N_2_) was added to the MEFs and incubated overnight (16 hours). MEFs were washed 3× in ice-cold PBS prepared with ultrapure water. MEFs were counted using a Denovix cell counter and 1mL of extraction buffer added/10^6^ cells. Extraction buffer (50% methanol, 30% acetonitrile, 20% ultrapure H_2_O, 50ng/mL HEPES made with LC-MS grade reagents). Cells were removed on ice and lysates were added to Eppendorf tubes and agitated for 15 minutes at 4°C and then incubated for 1 hour at -20°C. Lysates were centrifuged at 13,000rpm for 30 minutes at 4°C and the supernatant collected and added to autosampler tubes and stored at -80°C. LC-MS analysis was performed using a Q Exactive Hybrid Quadrupole-Orbitrap mass spectrometer coupled to a Dionex U3000 UHPLC system (Thermo Fisher Scientific). The liquid chromatography system was fitted with a Sequant ZIC-pHILIC column (150mm × 2.1mm) and guard column (20mm × 2.1mm) and temperature maintained at 45°C. The mobile phase was composed of 20mM ammonium carbonate and 0.1% ammonium hydroxide in water (solvent A) and acetonitrile (solvent B). The flow rate was set at 200nL/minute. The mass spectrometer was operated in full MS and polarity switching mode. The acquired spectra were analyzed using Xcalibur Qual Browser and Xcalibur Quan Browser software (Thermo Fisher Scientific). For metabolomic analysis using zebrafish, we used the protocol derived from ^35^. 50 fish (6 dpf) were placed in water and incubated for 24 hours in L-glucose U-^13^C_6_ [10mM],L-glutamine U-^13^C_5_ [5mM] and L-glutamine U-^15^N_2_ [5mM]. Fish were anaesthetized, snap-frozen and placed in Precellys homogenizer tubes with 500μl of extraction buffer and samples were processed as described.

### Proteomics

Tandem-mass-tagging (TMT) LC-MS proteomics was performed as described previously ^36^ using the King’s College London Proteomics Facility https://www.kcl.ac.uk/research/facilities/proteomics-facility. Each aliquoted sample was digested by following a routine in-solution digestion protocol prior to subsequent analysis by mass spectrometry (MS), including in-solution reduction, alkylation and digestion with trypsin. Cysteine residues were reduced with dithiothreitol and derivatized by treatment with iodoacetamide to form stable carbamidomethyl derivatives. The digestion was carried out with a mixture of trypsin and LysC overnight at room temperature after initial incubation at 37°C for 2 hours. The digested peptides were cleaned up with a spin column of PR-C18 resins. The cleaned peptides were resuspended in 100mM Triethylammonium bicarbonate for TMTpro tag labelling. TMTpro reagents (Thermo-Fisher Scientific; Lot #: XB341490/YB370079) were added to the peptides along with acetonitrile to achieve a final acetonitrile concentration of approximately 30% (v/v) in a total volume of 100µl. Following incubation at room temperature for 1 hour, the reaction was quenched with hydroxylamine to a final concentration of 0.3% (v/v), after labelling efficiency check. Finally, 16plex tags were combined as one TMTpro set and dried using a vacuum centrifuge (Savant SPD131DDA SpeedVac, Thermo-Fisher Scientific) for the next step. In order to explore a depth of proteome, a fractionation of TMTpro labelled peptide mix prior to liquid chromatography-MS (LC-MS) analysis was performed using Pierce High pH Reversed-Phase Peptide Fractionation Kit. In brief, the TMT labelled peptide mixture was fractionated into a total of 8 fractions. Peptide fractions were dried using a vacuum centrifuge and resuspended in 2 % acetonitrile, 0.05 % trifluoroacetic acid (in ultrapure water) and ready for LC-MS analysis. At this point, each TMTpro set generated eight fractions labelled from F1 to F8. Chromatographic separation was performed using an Ultimate 3000 NanoLC system (Thermo Fisher Scientific). Peptides were resolved by reversed phase chromatography on a 75µm*50cm C18 column using a three-step gradient of water in 0.1% formic acid (A) and 80% acetonitrile in 0.1% formic acid (B). The gradient was delivered to elute the peptides at a flow rate of 250nl/minute over 100 minutes. The eluate was ionized by electrospray ionization using an Orbitrap Fusion Lumos (Thermo-Fisher Scientific, UK) operating under Xcalibur v4.4. The instrument was programmed to acquire MS data using “Synchronous Precursor Selection with Multi-notch MS3” method by defining a 3s cycle time among a full MS scan, MS/MS fragmentation and MS3 fragmentation. Data were acquired using one full-scan MS spectrum at a resolution of 120,000 at 200m/z with 100% normalized AGC target and a scan range of 400∼1500m/z and maximum injection time of 100ms. The MS/MS fragmentation was conducted using collision-induced dissociation and quadrupole ion trap analyser. Parameters were set up as 100% normalized AGC target, NCE (normalized collision energy) 35, q-value 0.25, isolation window 1.2 Th and maximum injection time 50 ms. The MS3 scan was analyzed using higher-energy collision dissociation and Orbitrap analyser with a synchronous precursor selection. Parameters include number of SPS 10, NCE 65, 400% normalized AGC target, maximum injection time 105 ms, resolution 50,000 at 200 Th, isolation window 1.6. 2-6 charged states are defined within this method. After LC-MS acquisition, 8 raw files from one TMTpro set were generated and used for further analysis. Raw mass spectrometry data were processed into peak list files using Proteome Discoverer (Thermo-Fisher Scientific; v2.5) (PD 2.5). Processed data were then searched using Sequest search engine embedded in PD 2.5, against the current version of the reviewed Swissprot mouse databases downloaded from Uniprot (http://www.uniprot.org/uniprot/).

### Electron microscopy

Cells were fixed with 2.5% glutaraldehyde buffered with 100mM sodium cacodylate (pH7.2) for 24 hours at room temperature and stored for up to a week at 4°C. Sample processing and imaging were performed by the Histopathology Department (Great Ormond Street Hospital, London). Briefly, after secondary fixation in 1% osmium tetroxide (Agar Scientific) for 2 hours, samples were dehydrated in graded ethanols, transferred to propylene oxide then infiltrated and embedded in Agar 100 epoxy resin. Polymerization was carried out at 60°C for 48 hours. Ultrathin sections (90nm) were cut using a Diatome diamond knife on a Leica EM UC7 ultramicrotome. Sections were collected on copper grids and stained with alcoholic uranyl acetate and lead citrate. Sections were examined with a JEOL 1400 TEM and digital images recorded using an AMT XR80 digital camera. Approximately 50 images of WT and *Aip* KO MEFs and ∼50 images of healthy control fibroblasts and AIPd-PDFs taken. Autophagosomes and lysosomes were determined using the method previously described^37^.

### Lentivirus transduction

Aip was cloned into pLOC RFP IRES GFP (Dharmacon). This plasmid was transfected into HEK293T cells with packaging plasmids using a second-generation lentiviral system. Lentivirus was collected on day 4 and day 5 post transduction and concentrated using ultrafiltration columns (Sartorius). *Aip* KO MEFs were transduced with virus and GFP positive cells were isolated by cell sorting.

### Western blotting

Whole cell lysates were prepared following treatment. Cells were washed 3× with PBS and lysates generated using Phospho-Safe extraction buffer (Millipore) with complete protease inhibitors (Roche) or Ripa lysis buffer and incubated for 1 hour on a rotating wheel at 4°C. Samples were centrifuged at 17,000rpm for 30 minutes at 4°C, and supernatants quantitated using Bradford protein assay kit (Biorad). To examine ubiquitylated proteins, the lysis buffer was mixed with DUB inhibitors N-Ethylmaleimide (NEM) 10mM and PR-619 50µM. 6’ Laemmli SDS loading buffer was added and samples boiled at 95°C for 10 minutes, then run on a NuPAGE 4-12% Bis Tris gel in MES SDS running buffer (Invitrogen). Proteins were transferred using semi-dry transfer either to nitrocellulose blotting membrane 0.45µm (Amersham Protran) or PVDF membrane 0.45µm (Immobilon-FL) for total and phospho-proteins, respectively. Membranes were blocked in either 5% milk powder or 5% BSA for total and phospho-proteins. Antibodies were used at a dilution of 1:1,000 for immunoblotting and incubated overnight at 4°C. The following day, membranes were incubated with the corresponding IRDye secondary antibodies (LICOR) for 1 hour. Proteins were detected using the Odyssey Infrared Imaging System. Band densitometry was performed using the free graphical analysis software ImageJ (National Institutes of Health, Bethesda, Maryland, US).

### Confocal microscopy

Cells (50,000) were added to coverslips and the next day used for experiments. Cells were fixed with 4% PFA for 15 minutes and washed 3 × using PBS. Cells were permeabilized with 0.1% Triton X-100, 20µM digitonin, 0.5% saponin or methanol depending upon the target of interest. Coverslips were blocked with 5% BSA for 45 minutes before incubating the cells with primary antibodies in 1% BSA for 45 minutes. Coverslips were washed 3× and secondary antibodies in 1% BSA added and incubated for 45 minutes. Cells were counterstained with DAPI-mounting solution and placed on slides. Images were taken using a LSM880 (Zeiss) with a 63× objective. After acquisition, images were processed using ZEN software provided by Zeiss. Images were quantified using Image-J Software with the mean of 3-6 separate images (30-60 cells) used with the same threshold across all samples.

### Staining phosphoinositides

PI3P and PI4P were stained using a protocol previously described^27^. Briefly, cells were permeabilized for 5 minutes with 20µM digitonin in buffer A (20mM Pipes, pH 6.8, 137mM NaCl, and 2.7mM KCl). Cells were then washed 3× in buffer A, and then blocked for 45 minutes in buffer A containing 5% BSA. PI3P (GST-2×FYVE) (10µg) was added in buffer A for 60 minutes at room temperature. Cells were washed 3× with buffer A, anti-GST 488 in buffer A added to cells and incubated for 45-60 minutes. Cells were post-fixed for 5 minutes with 2% PFA, washed 3× with PBS containing 50mM NH_4_Cl, then once with MilliQ water and mounted.

### RNA sequencing

The primary analysis pipeline for RNAseq consisted of FastQC version 0.11.9 to look at the quality control within the reads data. HISAT2 version 2.2.1 ^38^ is used for read alignment. Sam Tools version 1.10 was used to convert SAM files to sorted and indexed bam files and Picard Metrics version 2.26.6 for further quality control. Quantification of the RNAseq data as counts per gene was completed using subread feature Counts version 2.01 ^39^. R studio (R version 4.2.2) was used for differential gene expression analysis using DESeq2 version 1.38.3^40^.

### Generation of aip loss of function zebrafish

To generate *aip* knock-out zebrafish, a crRNA was designed targeting *aip* exon 2 using CHOPCHOP (https://chopchop.cbu.uib.no), where the target site was included in the recognition site of BstX1 restriction enzyme [**<u>CCA</u>**<u>GGACGATGG</u>GAGGTCACAGT (PAM sequence in bold, restriction site underlined)]. 1nL of a solution containing 62.5 ng/ml crRNA, 62.5ng/ml tracrRNA and 5mM Cas9 (M0386M, NEB Ltd), was injected in one-cell stage zebrafish embryos (wild-type, TU). Around 100 embryos were injected and the knockout efficiency was assessed by PCR from genomic DNA (*aip*_Forward, 5’-TCATTACCGCACTAGCCTGT-3’; *aip*_Reverse, 5’-TGAATTCAGCTATTTCCCCCTCTT-3’) followed by BstX1 restriction enzyme digestion and 3% gel electrophoresis. The higher percentage of non-digested PCR products indicated higher efficiency and lower mosaic rate. Around 50 injected fish were raised to adulthood and crossed with wild-type TU fish to identify founders. To genotype, either the whole larva was collected individually in a 96-well PCR plate, or adult fish were anaesthetized with MS-222 at 168mg/l (E10521, NEB Ltd) and a caudal fin clipping was collected for genomic DNA extraction and amplified by PCR primers mentioned above. The amplicons were sequenced at Source Bioscience, UK. The molecular and morphological experiments were conducted among the third generation (F3) including *aip* wild-type, heterozygous and homozygous fish.

### Zebrafish maintenance

Zebrafish were housed in a recirculating system (Tecniplast, UK) on a 14-hour/10-hour light/dark cycle and a constant temperature of 28°C. Fish were fed twice daily with ZM-400 fry food (Zebrafish Management Ltd., UK) in the morning, and brine shrimp in the afternoon. Breeding was set up in the evening, either in sloping breeding tanks (Tecniplast, UK) or in tanks equipped with a container with marbles to isolate eggs from progenitors. For experiments where the developmental stage of larvae was important, we placed barriers between the fish to keep them isolated in the breeding tank. The following morning, barriers were removed to allow spawning. Eggs were collected in 90mm Petri dishes the following morning, sorted fertile from infertile, and then incubated at 28°C (maximum 50 eggs/dish). Dishes were checked daily to ensure consistent developmental stage across groups. If reared, larvae were moved to the recirculating system at 5 days post fertilization (dpf). All procedures were carried out under license (CB license number: PP6385208) in accordance with the Animals (Scientific Procedures) Act, 1986 and under guidance from the Local Animal Welfare and Ethical Review Board at Queen Mary University of London.

### Zebrafish morphological analysis

From 5 dpf all larvae were assessed twice daily for effects on morphology or behavior. Any larvae showing altered morphology such as yolk sac oedema, body curvature, altered swimming or lack of movement were killed in accordance with UK animal legislation. For size measurements, zebrafish larvae were anaesthetized using MS-222 at 168mg/l and mounted in 3% methyl cellulose in E3 embryo medium. Images were taken by a Leica S9i stereo microscope with an integrated 10 M pixels network camera. The body length was measured using the free graphical analysis software ImageJ (National Institutes of Health, Bethesda, Maryland, US). Following imaging, larvae were washed to remove methyl cellulose and transferred individually to single wells of a multi-well plate or single tanks for genotyping.

Zebrafish larvae were anaesthetized using MS-222 at 168mg/l and mounted in 3% methyl cellulose in E3 embryo medium. Images were taken by a Leica S9i stereo microscope with an integrated 10 M pixels network camera. The body length was measured using ImageJ. Following imaging, larvae were washed to remove methyl cellulose and transferred individually to single wells of a multi-well plate for genotyping.

### Genotyping of larval fish

For experiments where no further analysis was necessary after genotyping (morphological analysis and survival experiments), genomic DNA was extracted from the whole larvae with 100μL of 50mM sodium hydroxide before incubation at 95°C for 30 minutes, followed by the addition of 10μl of 1mM Tris-HCl. For immunohistochemical analysis, individual larvae were fixed with 4% paraformaldehyde, and then only tail sections were collected and incubated in 25μL of 50mM NaOH at 95°C for 20 minutes, followed by the addition of 2.5μl of 1mM Tris-HCl. The genotyping PCR primers and procedure were as described above.

### Zebrafish immunofluorescence

Fixed and genotyped larvae were washed briefly with PBS. Samples were cryoprotected in 15% and 30% sucrose overnight at 4°C before being embedded in optimal cutting temperature compound in a cold bath with dry ice. Sectioning of blocks at 12μm thickness was performed using a cryostat and mounted on SuperFrost Plus microscope slides (Thermo-Fisher Scientific). Primary antibodies including Anti-Ubiquitin 1:250 and anti-LC3B 1:200 and goat anti-rabbit secondary antibody Alexa Fluor™ 546 were used for immunofluorescence following standard protocol ^41^. All slides were counterstained with DAPI (1μg/ml) for 10 minutes.

### Assessing autophagy in larval zebrafish

The induction of autophagy was assessed following acute dietary restriction. Zebrafish larvae were reared under normal conditions until 5 dpf. At 5 dpf WT and homozygous mutant larvae were separated into 2 groups for normal and restricted feeding. The normal group was fed normally twice a day with paramecium from 5-7 dpf while the restricted group were fed normally at 5 dpf but received no further paramecium before being harvested for protein analysis.

### Zebrafish dietary restriction conditions

Larvae were randomly divided into two groups at 5 dpf and each group included six biological replicates with 25 embryos each. The control group was fed with paramecium twice a day. The dietary restricted group was fed normally on day 5 but received no further paramecium. All other rearing conditions were as normal. Larvae were checked twice a day for signs of harm, such as yolk sac oedema, body curvature, and lack of movement even upon stimulus. Any larvae showing these symptoms were humanely euthanized and recorded as dead. After euthanasia, the entire fish was collected for genomic DNA extraction and genotyping. Larvae were reared until 11 dpf and number of larvae removed each day recorded.

### Zebrafish Western blotting

Zebrafish larvae were reared under normal fed conditions until 5 dpf and underwent dietary restriction for 12 hours and harvested for Western blot analysis Zebrafish were put in 200μl cold RIPA lysis buffer with 1mM PMSF. Samples were then homogenized using a bead mill homogenizer and incubated on a shaker for 30 minutes at 4°C before centrifuging at 13,000rpm, 4°C for 15 minutes and Western blot analysis was performed. NIH software Image J was applied for blot scanning and protein area quantifications.

### Zebrafish Proteasome activity

Proteasome activity was analyzed using Proteasome Activity Assay Kit. Groups of wild-type and homozygous larvae were generated as described above for metabolomic analysis. 10 fish from each group were homogenized in 200µl 0.5% NP-40 in distilled water by pipetting up and down and then centrifuged for 15 minutes at 13,500 rpm, 4°C and the assay was performed according to the manufacturer’s instructions.

### Zebrafish LysoTracker staining

LysoTracker staining was performed as previously reported ^42^. Larvae were incubated in 5µM LysoTracker Deep Red in E3 medium for 2 hours with gentle shaking and washed 3× with E3 medium. Live images were acquired with Leica fluorescence stereomicroscope MZFLIII (Leica).

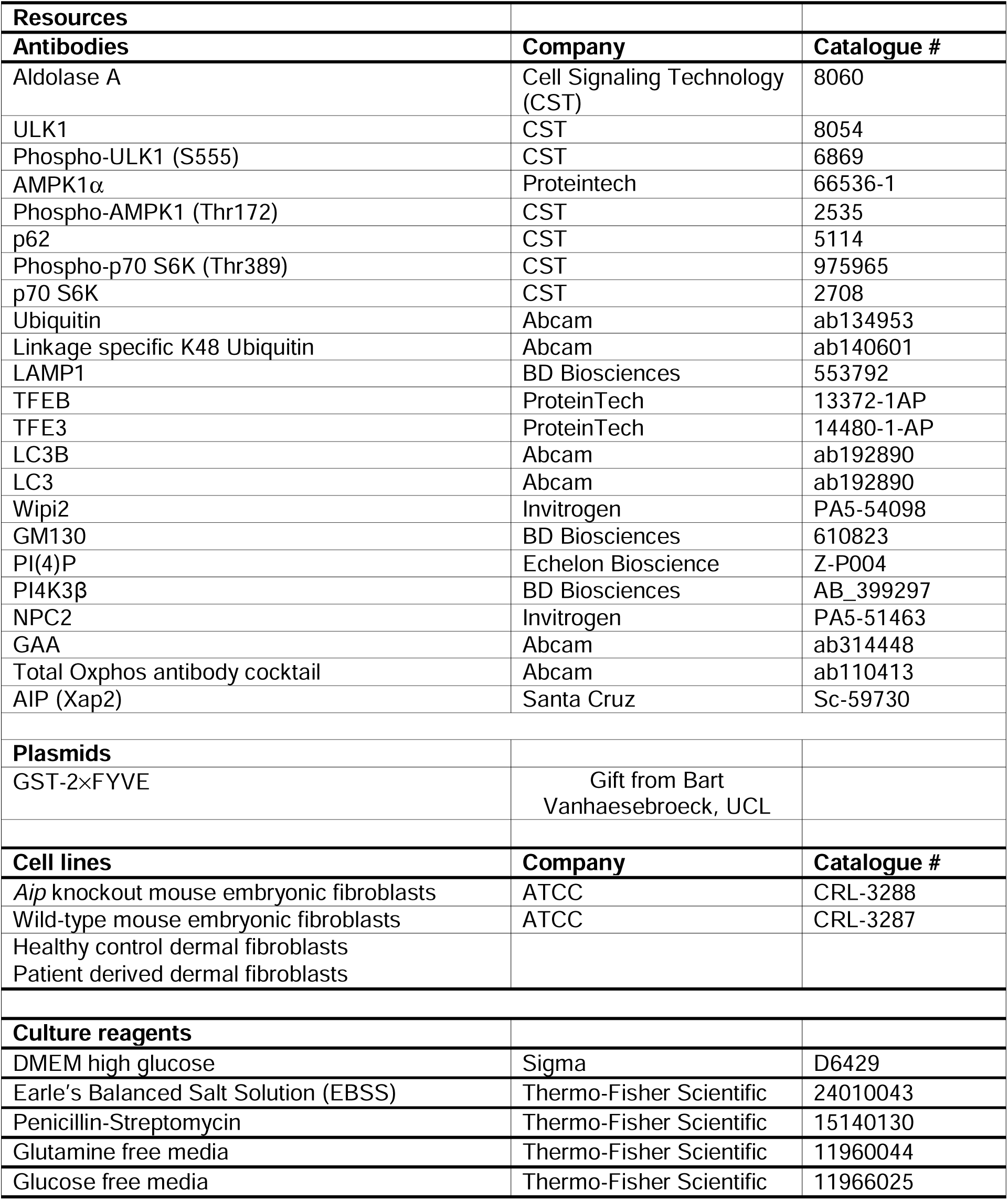
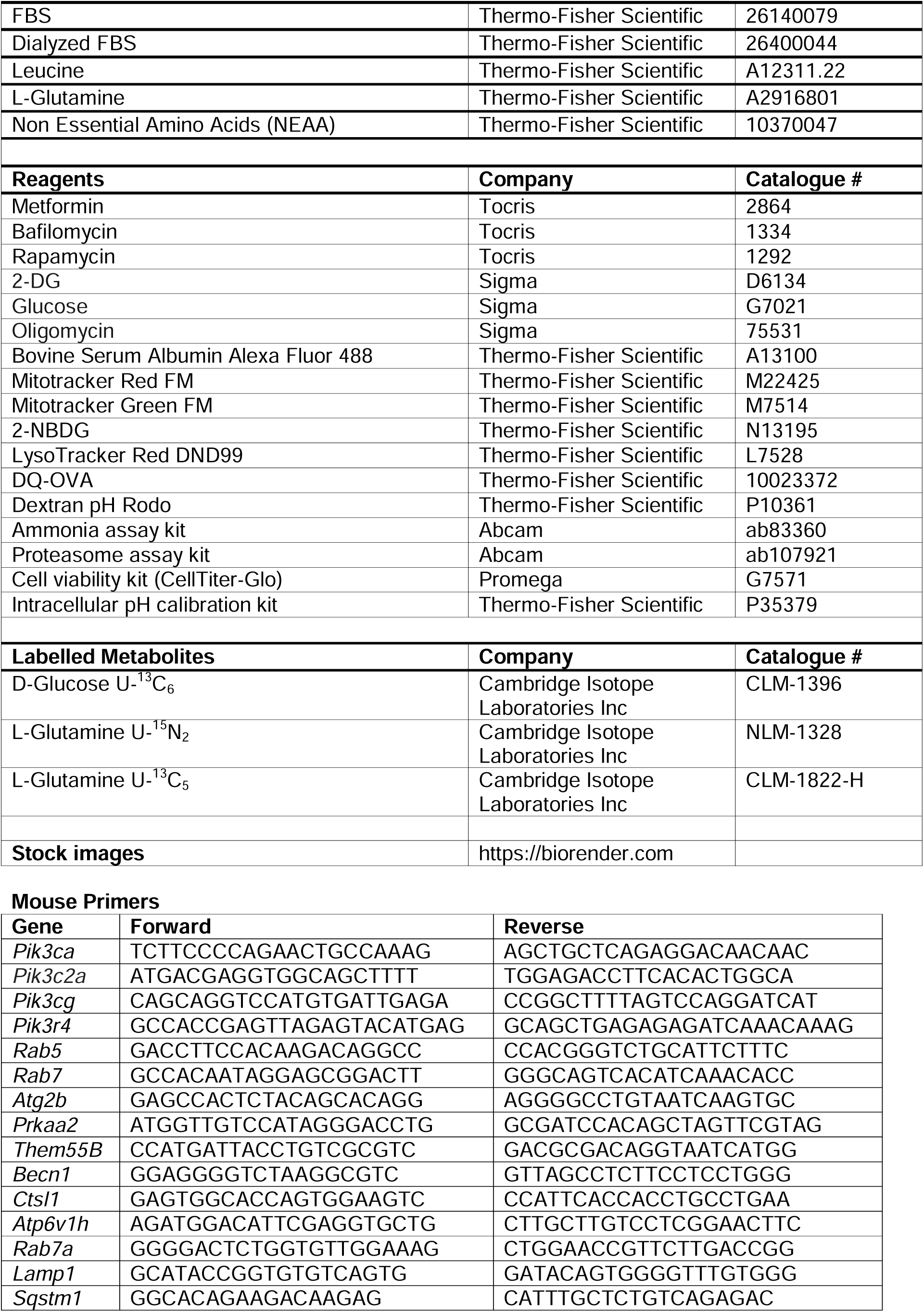
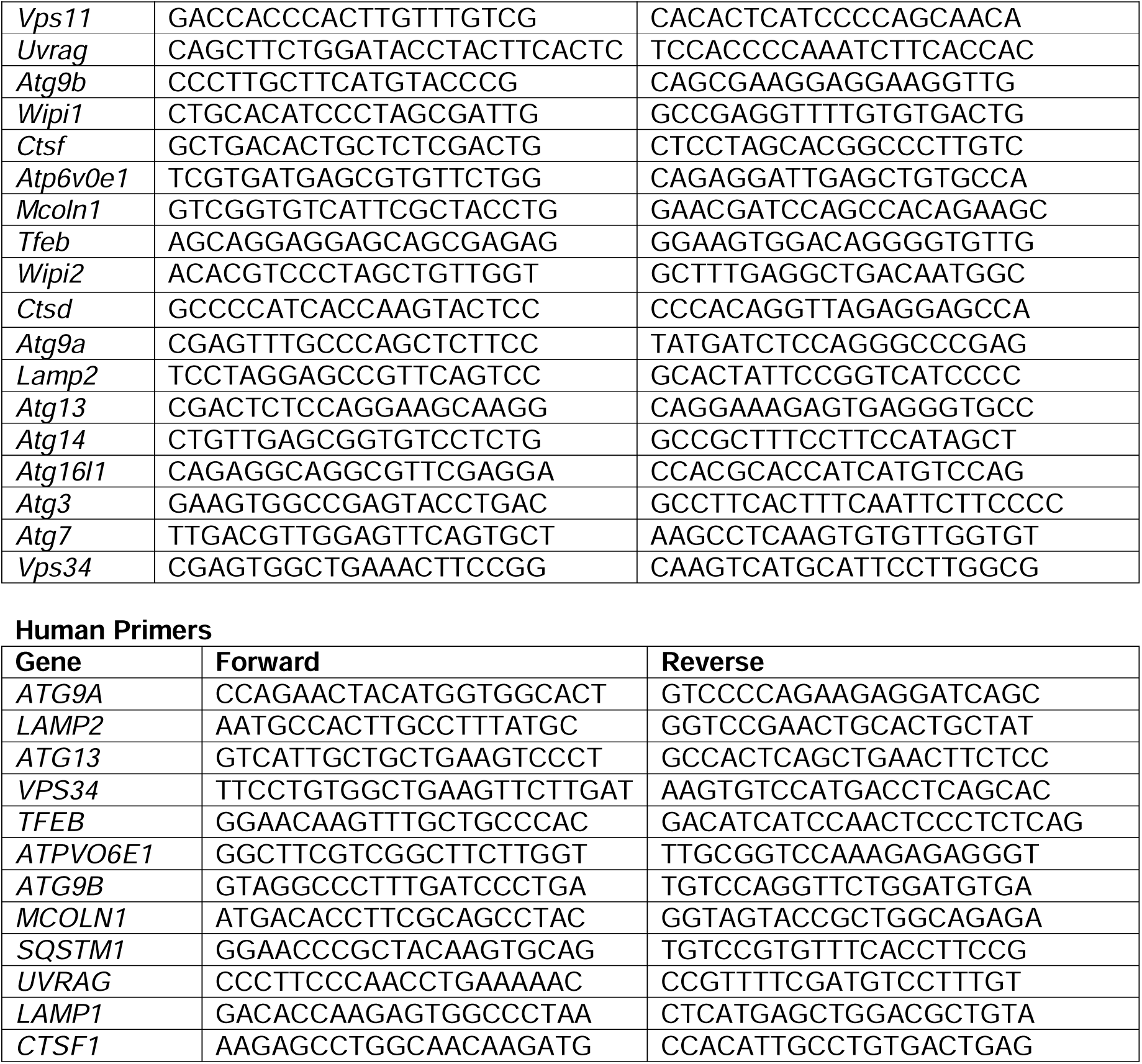

### Statistical analysis

Statistics were performed using Graphpad Prism version 9.2 software. Comparisons were made with Student’s *t*-test and one-way or two-way ANOVA with Tukey’s multiple comparison test. Significance was taken as p<0.05 and marked on figures as *p <0.05, **p <0.01, ***p <0.001, ****p <0.0001. Horizontal bars represent the mean values. Error bars on figures represent standard error (SEM).

## Ethics

Parents signed written consent to take part in the study and the study was approved by the Ethic Committee (MREC 06/Q0104/133).

CB Home Office license number: PP6385208

## Data and materials availability

MEFs and *aip* KO zebrafish used in this study are available upon request.

## Author contributions

**Conceptualization:** MK, OH

**Patient identification and care:** HTB, VÖE, KAR, HVE, VEK, AD, RZ, YS, AOB

**Investigation:** XW, SB, CTL, OS, NU, MLV, GA, SDT, SMH, VM, KB, MD, KS, CYC, GB, CB, YY, OH

**Bioinformatics and data analysis:** CH, EA, JPC, LP, ST, SP, OH

**Funding acquisition:** MK, CB, OH

**Writing:** MK, JPC, XW, CB, GC, EA, OH

## Funding

Rosetrees Trust [M789] (OH, MK)

Great Ormond Street Hospital Children’s Charity (registered charity no.1160024) [V4722]) (OH, MK)

China Scholarship Council PhD scholarship (XW, MK, CB)

Clinical Training Fellowship from The Medical College of St Bartholomew’s Trust [MEAG1T1R] (CTL, MK)

Medical Research Council [MR/M018539/1] (MK) Barts Charity [MGU0549] (MK)

William Harvey Research Institute [MGU093] (JPC)

## Acknowledgments

We are grateful to the patients and their families for providing samples and data for the study. We acknowledge the help of Dr Steve Lynam and Dr. Xiaoping Yang from the Kings College London proteomics facility. We are grateful to Dr James Davison and Professor Detlef Bockenhauer at Great Ormond Street Hospital in London, UK and to numerous colleagues who have provided reagents and helpful discussions regarding this work.

---

**Figure S1.**
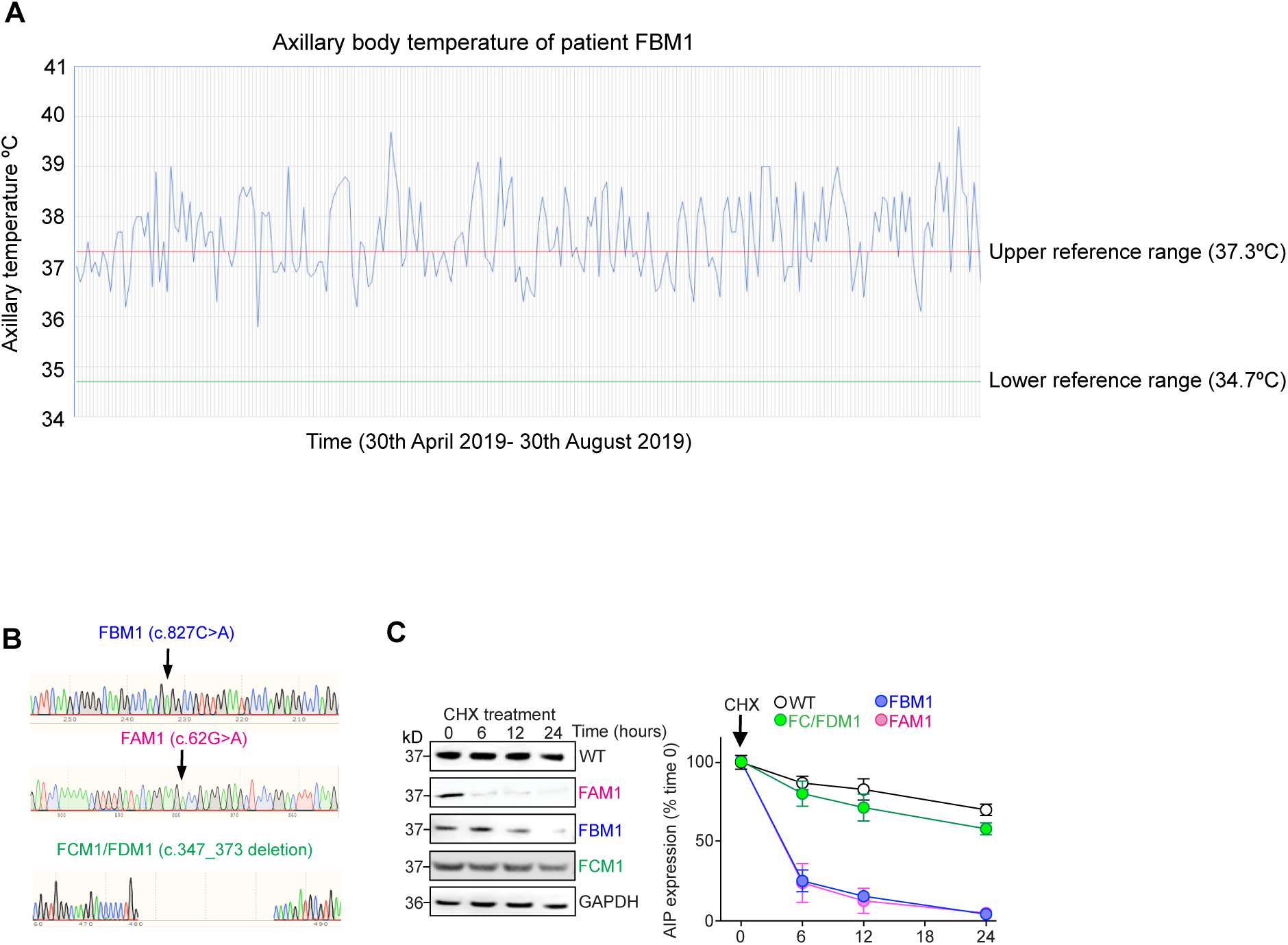

**Figure S2.**
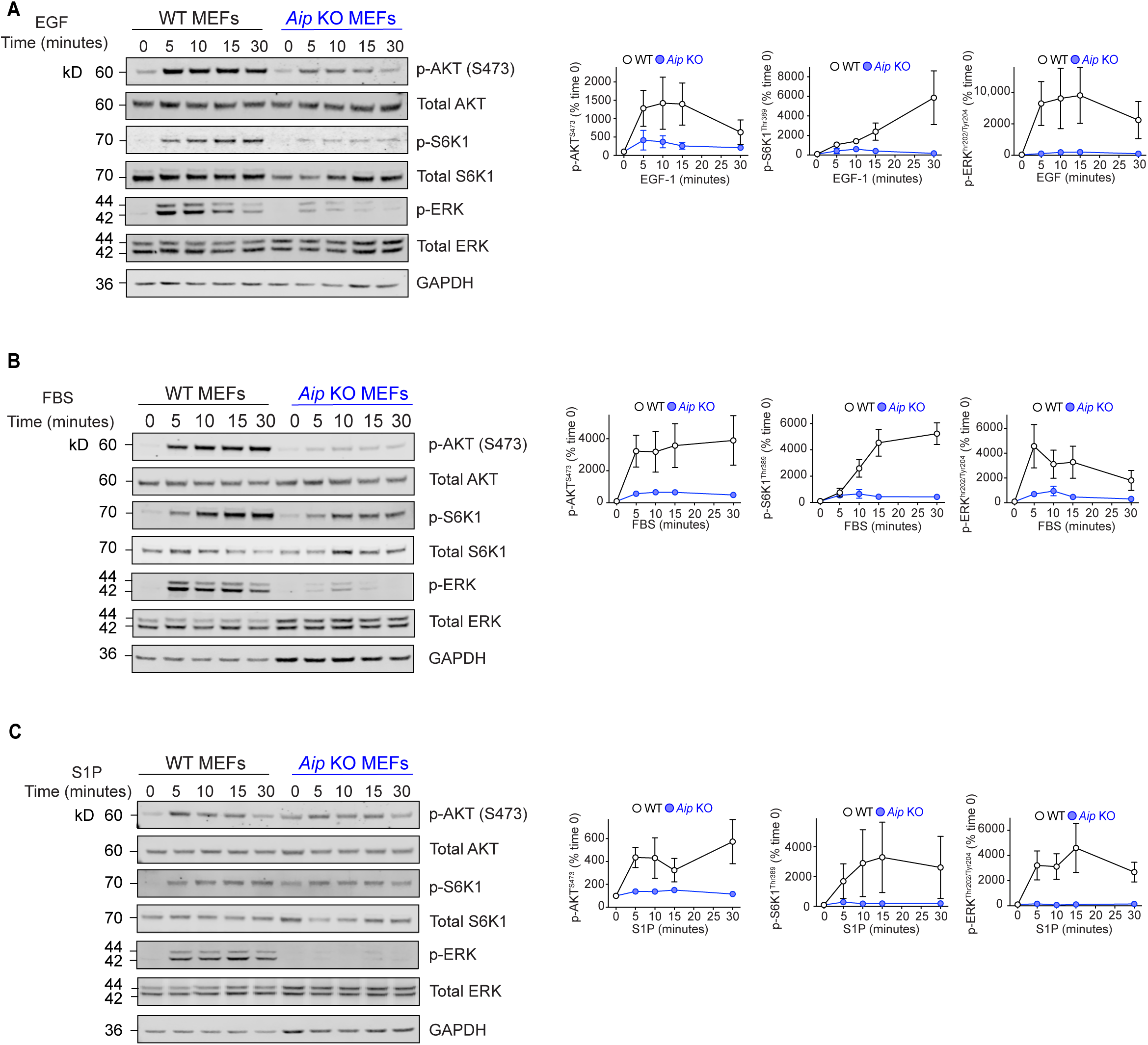

**Figure S3.**
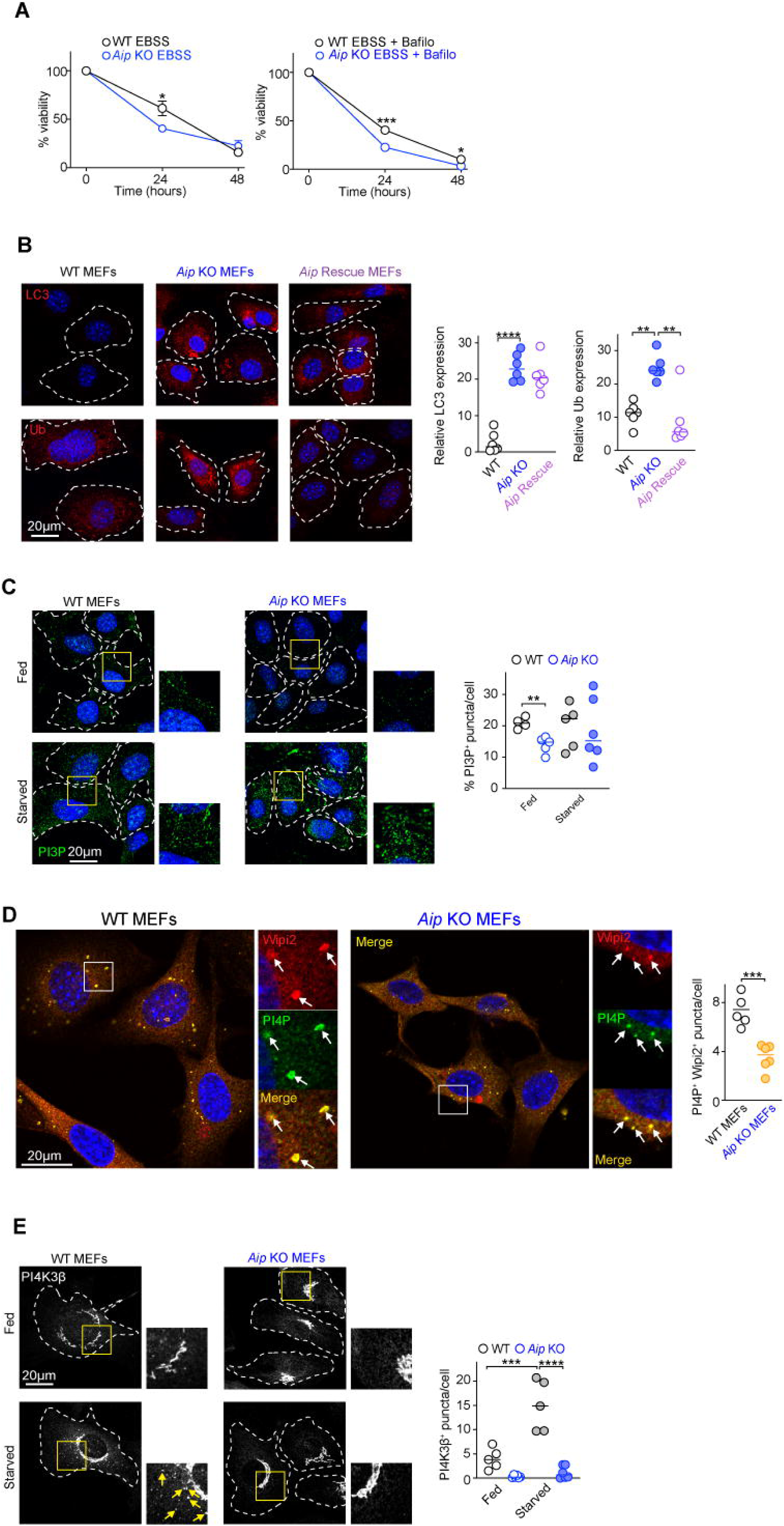

**Figure S4.**
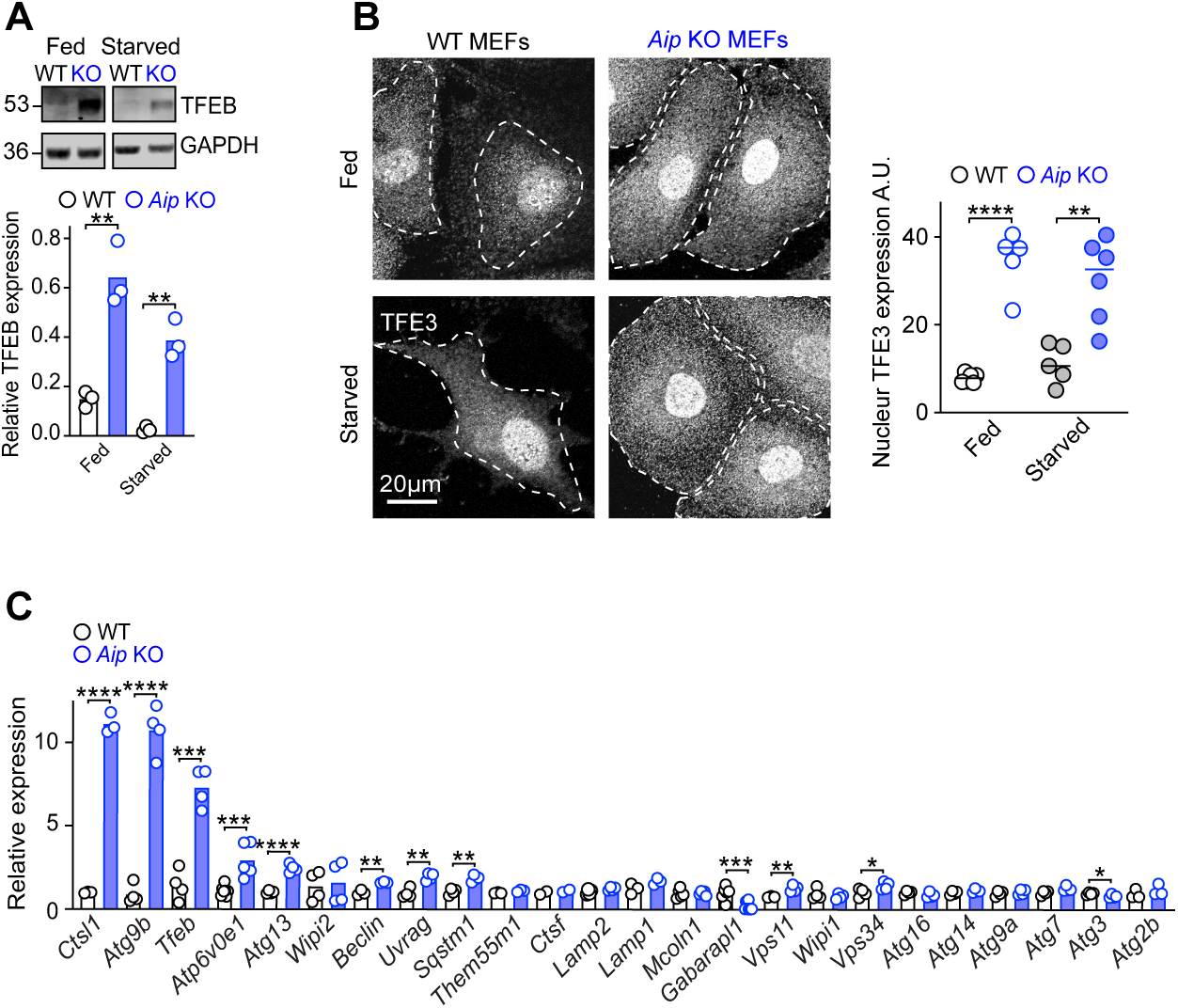

**Figure S5.**
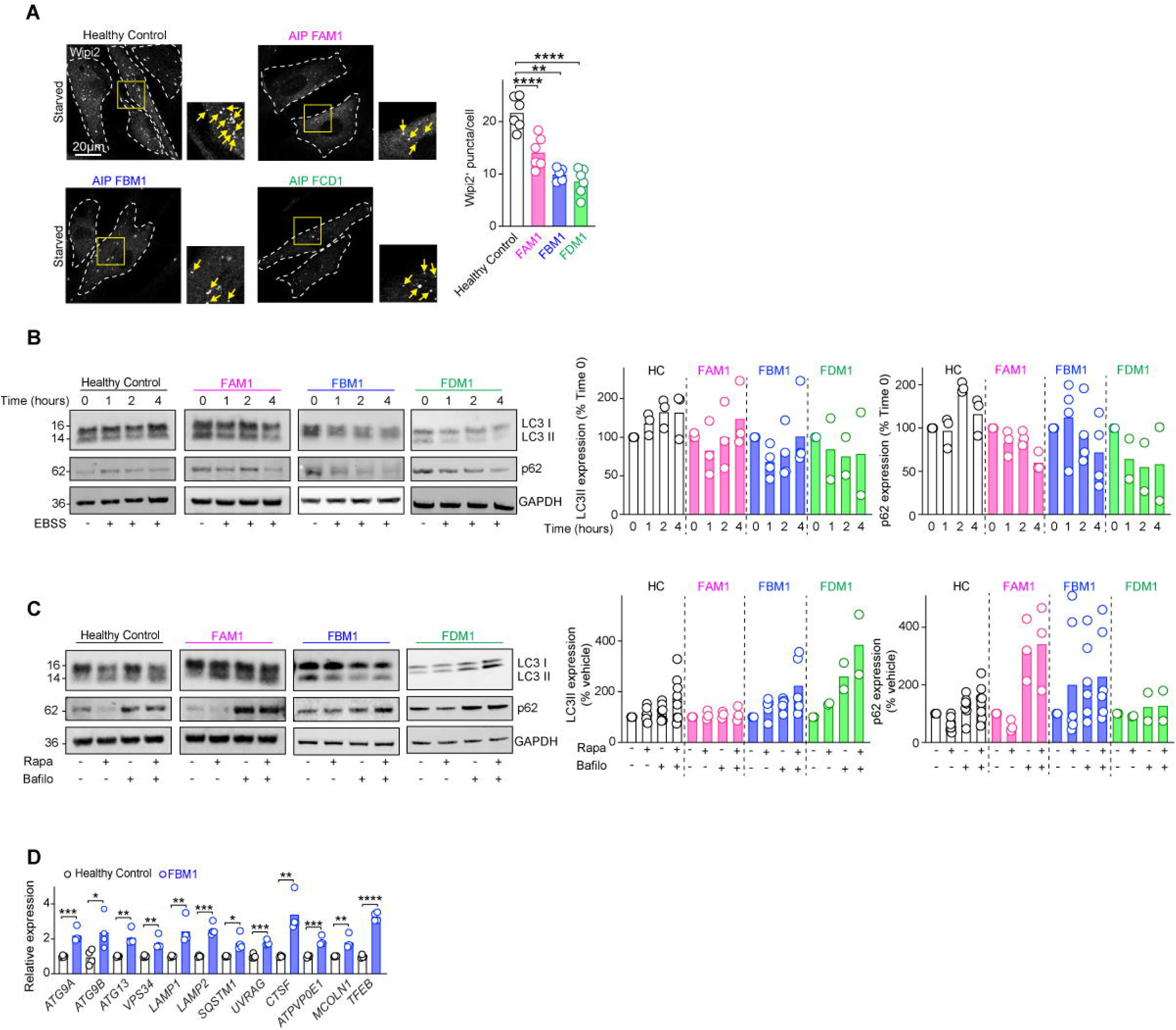

**Figure S6.**
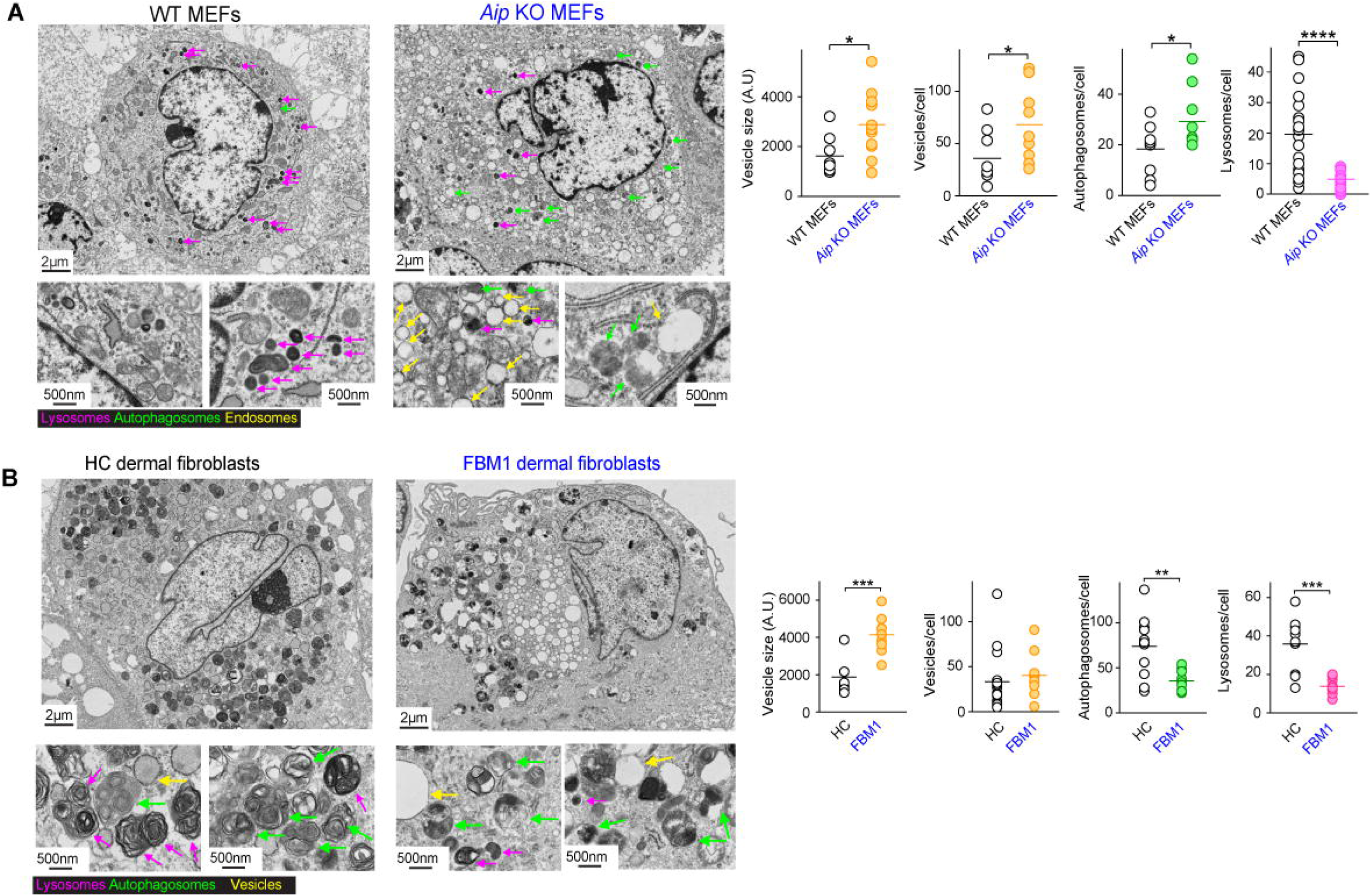

**Figure S7.**
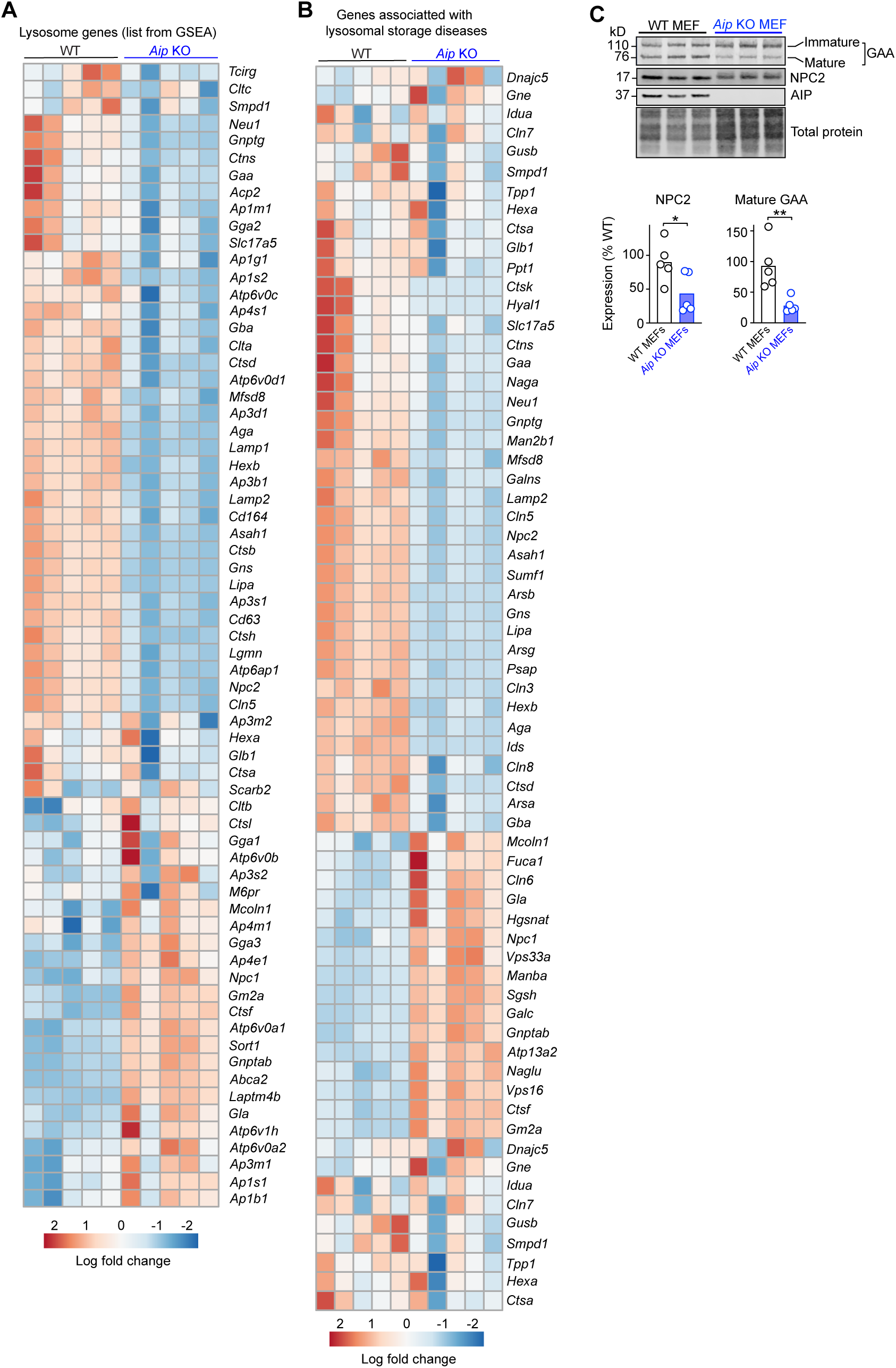

**Figure S8.**
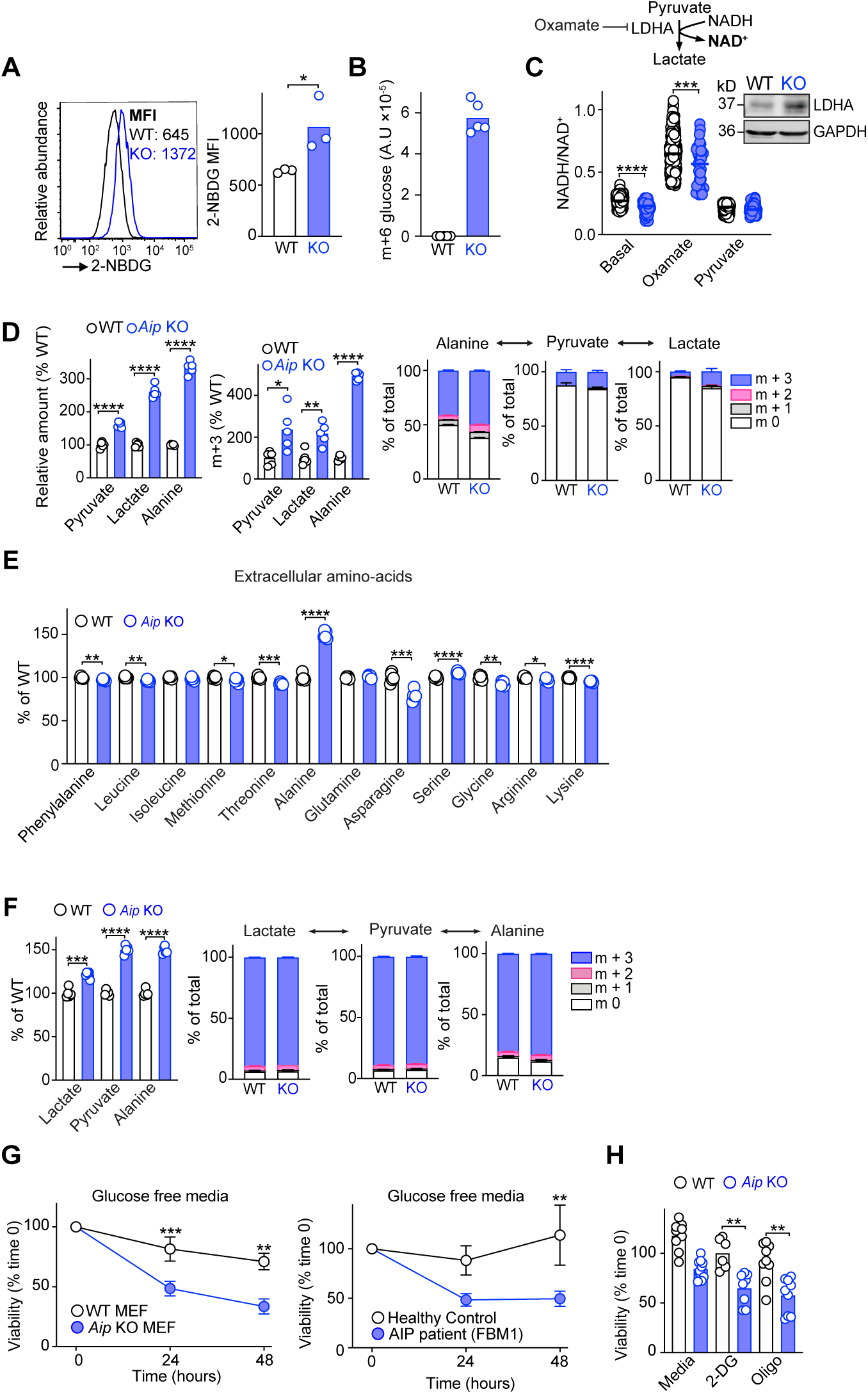

**Figure S9.**
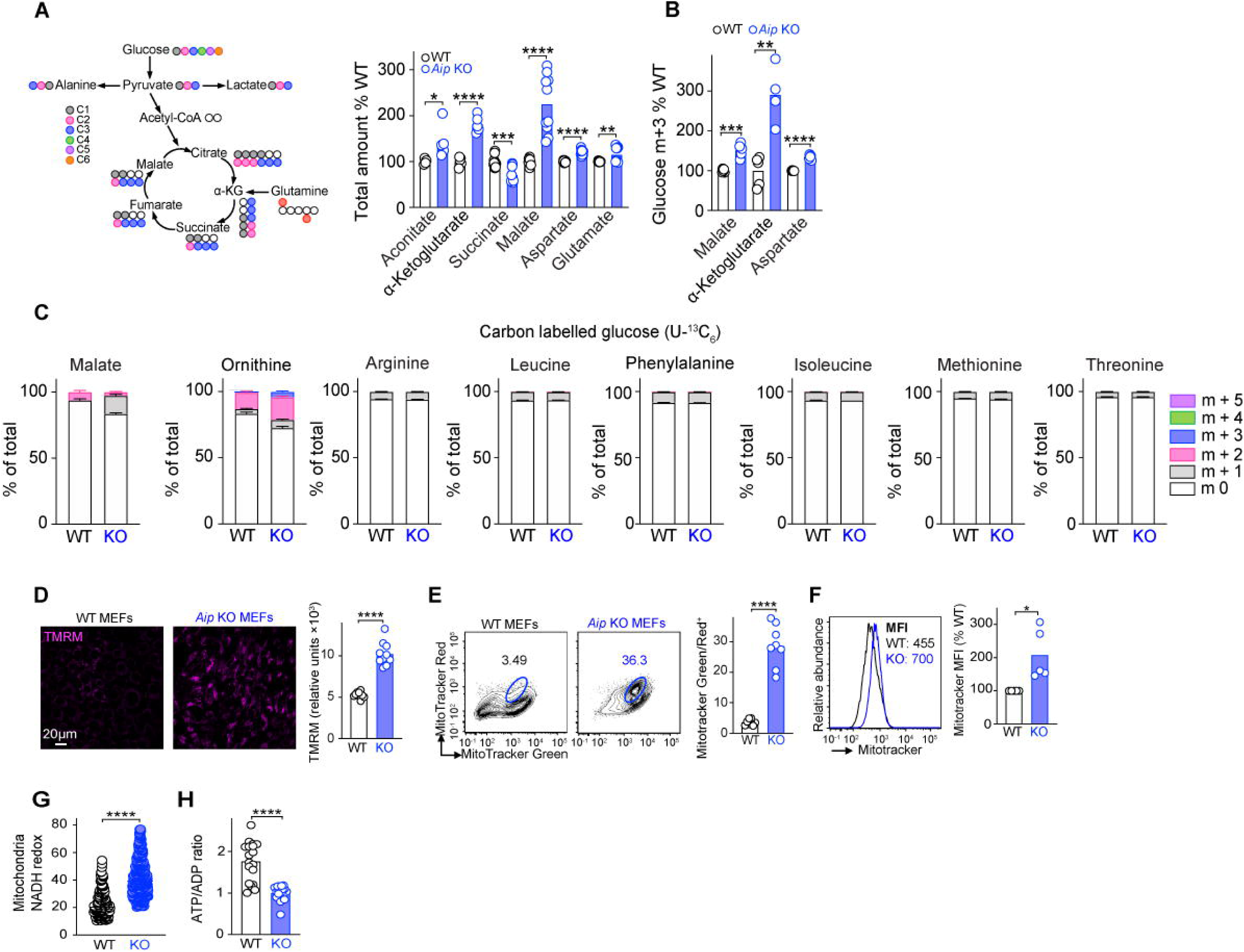

**Figure S10.**
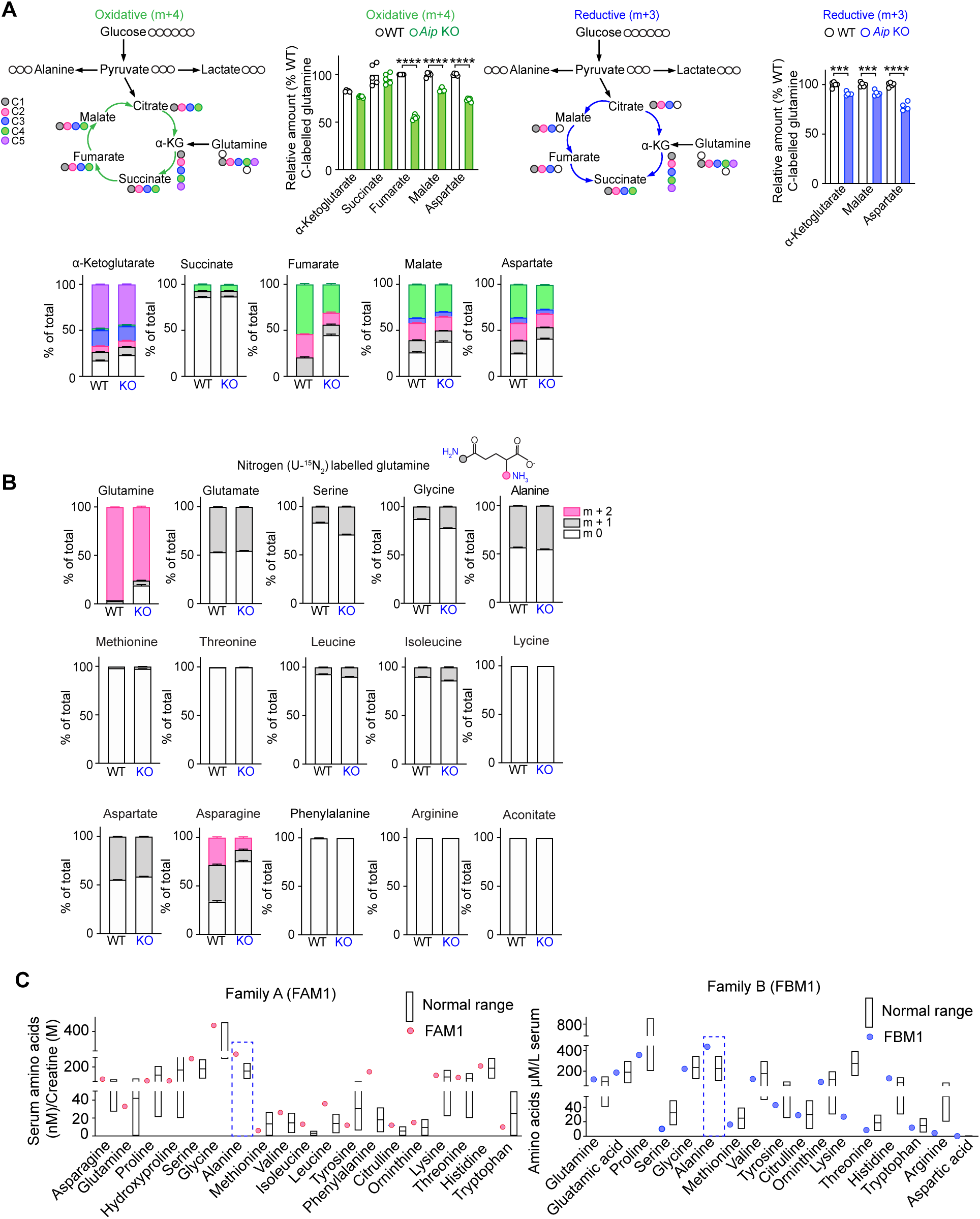

**Figure S11.**
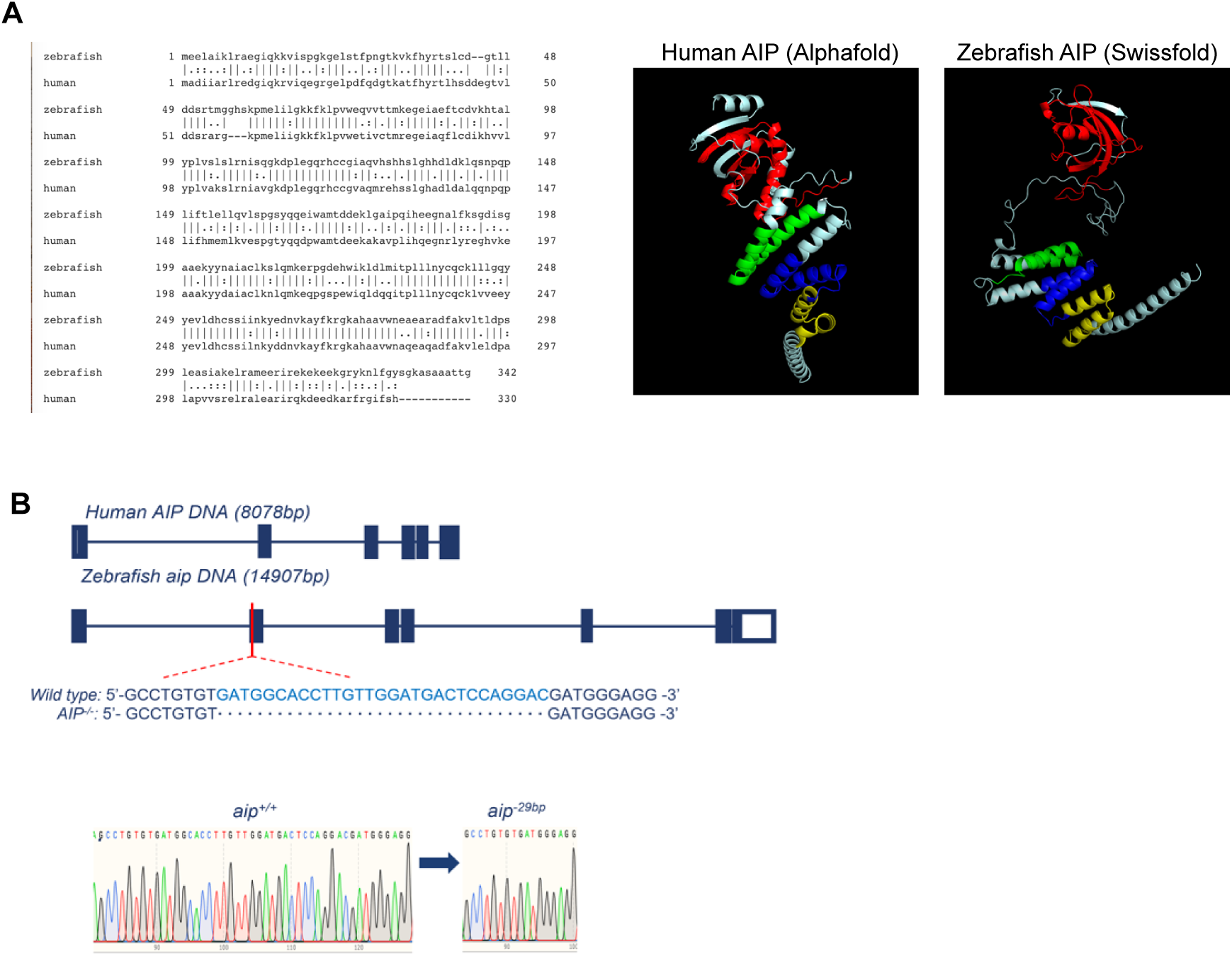

**Figure S12.**
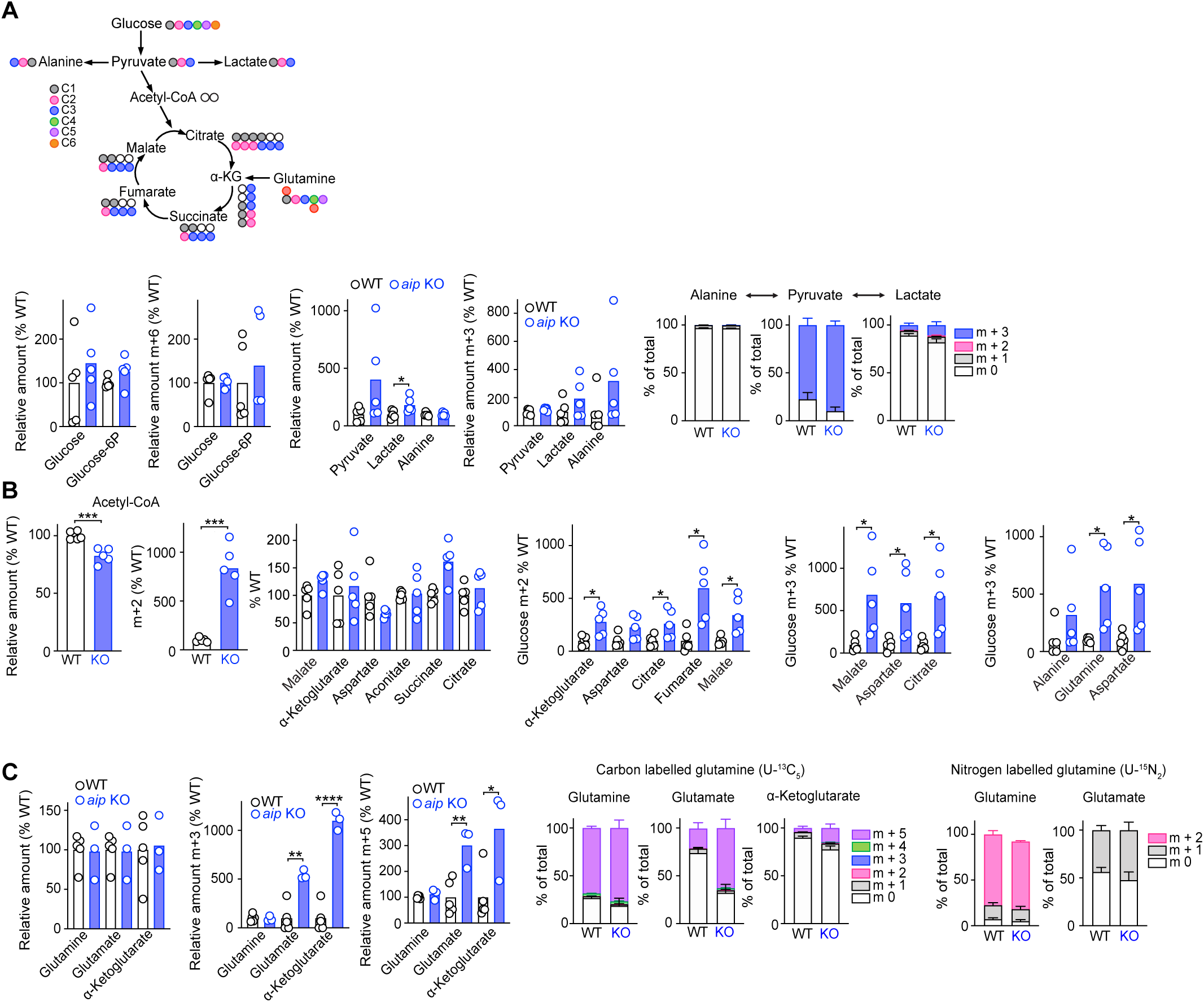

**Figure S13.**
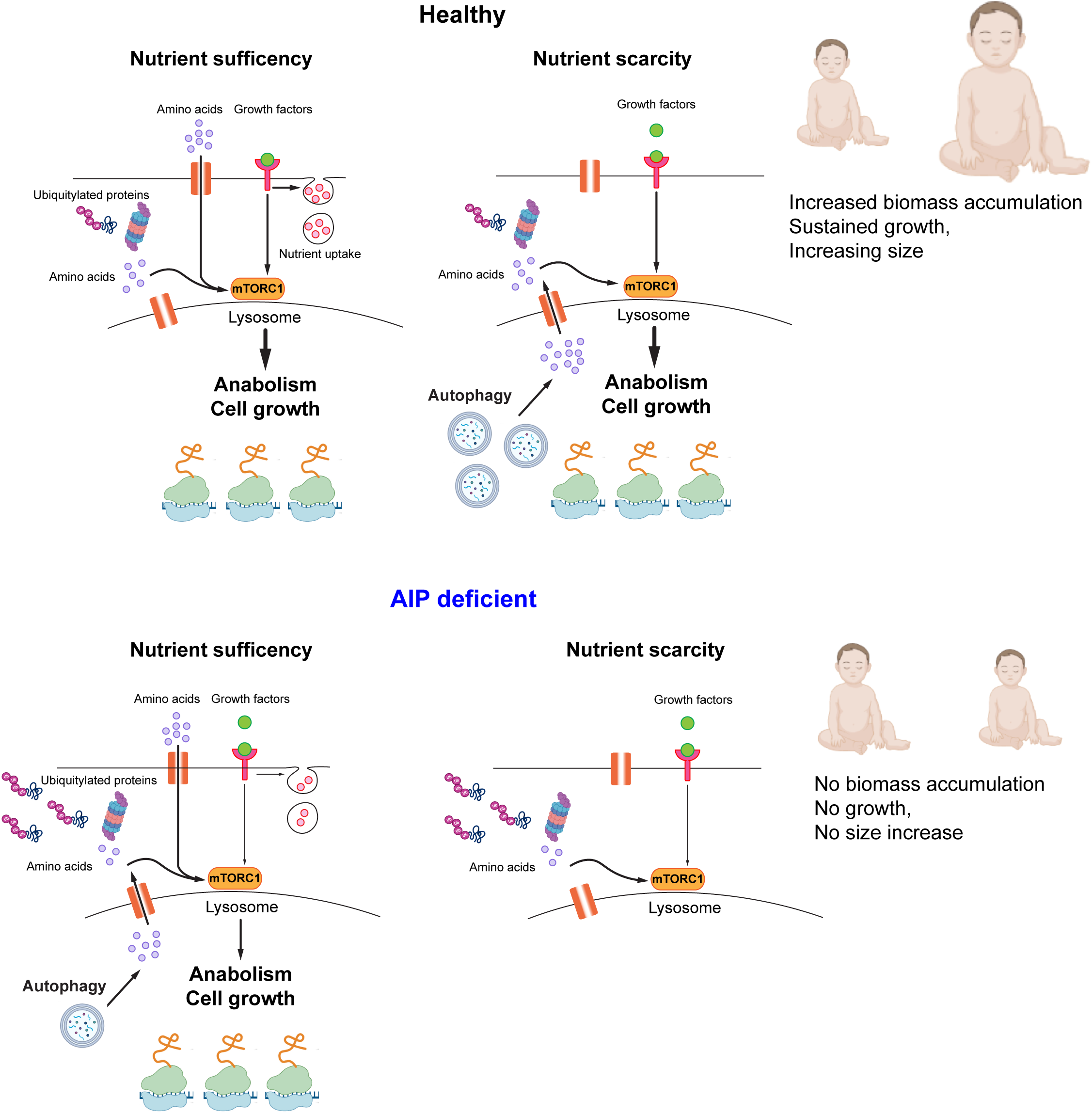

## REFERENCES

1. Laplante, M., and Sabatini, D.M. (2012). mTOR signaling in growth control and disease. Cell 149, 274–293. 10.1016/j.cell.2012.03.017.

2. Joshi, S., Wang, T., Araujo, T.L.S., Sharma, S., Brodsky, J.L., and Chiosis, G. (2018). Adapting to stress - chaperome networks in cancer. Nat Rev Cancer 18, 562–575. 10.1038/s41568-018-0020-9.

3. Schopf, F.H., Biebl, M.M., and Buchner, J. (2017). The HSP90 chaperone machinery. Nat Rev Mol Cell Biol 18, 345–360. 10.1038/nrm.2017.20.

4. Hartl, F.U., Bracher, A., and Hayer-Hartl, M. (2011). Molecular chaperones in protein folding and proteostasis. Nature 475, 324–332. 10.1038/nature10317.

5. Butte, N.F. (2005). Energy requirements of infants. Public Health Nutr 8, 953–967. 10.1079/phn2005790.

6. Tippetts, T.S., Sieber, M.H., and Solmonson, A. (2023). Beyond energy and growth: the role of metabolism in developmental signaling, cell behavior and diapause. Development 150. 10.1242/dev.201610.

7. Ward Platt, M., and Deshpande, S. (2005). Metabolic adaptation at birth. Semin Fetal Neonatal Med 10, 341–350. 10.1016/j.siny.2005.04.001.

8. Wolff, S., Weissman, J.S., and Dillin, A. (2014). Differential scales of protein quality control. Cell 157, 52–64. 10.1016/j.cell.2014.03.007.

9. Levine, B., and Kroemer, G. (2019). Biological Functions of Autophagy Genes: A Disease Perspective. Cell 176, 11–42. 10.1016/j.cell.2018.09.048.

10. Efeyan, A., Zoncu, R., Chang, S., Gumper, I., Snitkin, H., Wolfson, R.L., Kirak, O., Sabatini, D.D., and Sabatini, D.M. (2013). Regulation of mTORC1 by the Rag GTPases is necessary for neonatal autophagy and survival. Nature 493, 679–683. 10.1038/nature11745.

11. Kuma, A., Hatano, M., Matsui, M., Yamamoto, A., Nakaya, H., Yoshimori, T., Ohsumi, Y., Tokuhisa, T., and Mizushima, N. (2004). The role of autophagy during the early neonatal starvation period. Nature 432, 1032–1036. 10.1038/nature03029.

12. Komatsu, M., Waguri, S., Ueno, T., Iwata, J., Murata, S., Tanida, I., Ezaki, J., Mizushima, N., Ohsumi, Y., Uchiyama, Y., et al. (2005). Impairment of starvation-induced and constitutive autophagy in Atg7-deficient mice. J Cell Biol 169, 425–434. 10.1083/jcb.200412022.

13. Maweed Suzan Attia, Z.J., Ren Fan, Mei Jie (2021). *atg7* and *beclin1* are essential for energy metabolism and survival during the larval-to-juvenile transition stage of zebrafish. Aquaculture and Fisheries 7, (4) 359–72.

14. Goul, C., Peruzzo, R., and Zoncu, R. (2023). The molecular basis of nutrient sensing and signalling by mTORC1 in metabolism regulation and disease. Nat Rev Mol Cell Biol 24, 857–875. 10.1038/s41580-023-00641-8.

15. Collier, J.J., Guissart, C., Oláhová, M., Sasorith, S., Piron-Prunier, F., Suomi, F., Zhang, D., Martinez-Lopez, N., Leboucq, N., Bahr, A., et al. (2021). Developmental Consequences of Defective ATG7-Mediated Autophagy in Humans. N Engl J Med 384, 2406–2417. 10.1056/NEJMoa1915722.

16. Trivellin, G., and Korbonits, M. (2011). AIP and its interacting partners. J Endocrinol 210, 137–155. 10.1530/JOE-11-0054.

17. Vierimaa, O., Georgitsi, M., Lehtonen, R., Vahteristo, P., Kokko, A., Raitila, A., Tuppurainen, K., Ebeling, T.M., Salmela, P.I., Paschke, R., et al. (2006). Pituitary adenoma predisposition caused by germline mutations in the AIP gene. Science 312, 1228–1230. 10.1126/science.1126100.

18. Coopmans, E.C., and Korbonits, M. (2022). Molecular genetic testing in the management of pituitary disease. Clin Endocrinol (Oxf) 97, 424–435. 10.1111/cen.14706.

19. Sobreira, N., Schiettecatte, F., Valle, D., and Hamosh, A. (2015). GeneMatcher: a matching tool for connecting investigators with an interest in the same gene. Hum. Mutat. 36, 928–930. 10.1002/humu.22844.

20. Sun, D., Stopka-Farooqui, U., Barry, S., Aksoy, E., Parsonage, G., Vossenkämper, A., Capasso, M., Wan, X., Norris, S., Marshall, J.L., et al. (2019). Aryl Hydrocarbon Receptor Interacting Protein Maintains Germinal Center B Cells through Suppression of BCL6 Degradation. Cell Rep 27, 1461–1471.e1464. 10.1016/j.celrep.2019.04.014.

21. Kirkin, V., McEwan, D.G., Novak, I., and Dikic, I. (2009). A role for ubiquitin in selective autophagy. Mol Cell 34, 259–269. 10.1016/j.molcel.2009.04.026.

22. Suraweera, A., Munch, C., Hanssum, A., and Bertolotti, A. (2012). Failure of amino acid homeostasis causes cell death following proteasome inhibition. Mol Cell 48, 242–253. 10.1016/j.molcel.2012.08.003.

23. Korolchuk, V.I., Menzies, F.M., and Rubinsztein, D.C. (2010). Mechanisms of cross-talk between the ubiquitin-proteasome and autophagy-lysosome systems. FEBS Lett 584, 1393–1398. 10.1016/j.febslet.2009.12.047.

24. Gonzalez, A., Hall, M.N., Lin, S.C., and Hardie, D.G. (2020). AMPK and TOR: The Yin and Yang of Cellular Nutrient Sensing and Growth Control. Cell Metab 31, 472–492. 10.1016/j.cmet.2020.01.015.

25. Dooley, H.C., Razi, M., Polson, H.E., Girardin, S.E., Wilson, M.I., and Tooze, S.A. (2014). WIPI2 links LC3 conjugation with PI3P, autophagosome formation, and pathogen clearance by recruiting Atg12-5-16L1. Mol Cell 55, 238–252. 10.1016/j.molcel.2014.05.021.

26. McGrath, M.J., Eramo, M.J., Gurung, R., Sriratana, A., Gehrig, S.M., Lynch, G.S., Lourdes, S.R., Koentgen, F., Feeney, S.J., Lazarou, M., et al. (2021). Defective lysosome reformation during autophagy causes skeletal muscle disease. J Clin Invest 131. 10.1172/JCI135124.

27. Judith, D., Jefferies, H.B.J., Boeing, S., Frith, D., Snijders, A.P., and Tooze, S.A. (2019). ATG9A shapes the forming autophagosome through Arfaptin 2 and phosphatidylinositol 4-kinase IIIβ. J Cell Biol 218, 1634–1652. 10.1083/jcb.201901115.

28. Settembre, C., Di Malta, C., Polito, V.A., Garcia Arencibia, M., Vetrini, F., Erdin, S., Erdin, S.U., Huynh, T., Medina, D., Colella, P., et al. (2011). TFEB links autophagy to lysosomal biogenesis. Science 332, 1429–1433. 10.1126/science.1204592.

29. Platt, F.M., d’Azzo, A., Davidson, B.L., Neufeld, E.F., and Tifft, C.J. (2018). Lysosomal storage diseases. Nat Rev Dis Primers 4, 27. 10.1038/s41572-018-0025-4.

30. Ballabio, A., and Bonifacino, J.S. (2020). Lysosomes as dynamic regulators of cell and organismal homeostasis. Nat Rev Mol Cell Biol 21, 101–118. 10.1038/s41580-019-0185-4.

31. Yu, L., McPhee, C.K., Zheng, L., Mardones, G.A., Rong, Y., Peng, J., Mi, N., Zhao, Y., Liu, Z., Wan, F., et al. (2010). Termination of autophagy and reformation of lysosomes regulated by mTOR. Nature 465, 942–946. 10.1038/nature09076.

32. Warner, J.R. (1999). The economics of ribosome biosynthesis in yeast. Trends Biochem Sci 24, 437–440. 10.1016/s0968-0004(99)01460-7.

33. Hernández-Ramírez, L.C., Martucci, F., Morgan, R.M., Trivellin, G., Tilley, D., Ramos-Guajardo, N., Iacovazzo, D., D’Acquisto, F., Prodromou, C., and Korbonits, M. (2016). Rapid Proteasomal Degradation of Mutant Proteins Is the Primary Mechanism Leading to Tumorigenesis in Patients With Missense AIP Mutations. J Clin Endocrinol Metab 101, 3144–3154. 10.1210/jc.2016-1307.

34. Chung, C.Y., Singh, K., Kotiadis, V.N., Valdebenito, G.E., Ahn, J.H., Topley, E., Tan, J., Andrews, W.D., Bilanges, B., Pitceathly, R.D.S., et al. (2021). Constitutive activation of the PI3K-Akt-mTORC1 pathway sustains the m.3243CAC>CG mtDNA mutation. Nat Commun 12, 6409. 10.1038/s41467-021-26746-2.

35. Naser, F.J., Jackstadt, M.M., Fowle-Grider, R., Spalding, J.L., Cho, K., Stancliffe, E., Doonan, S.R., Kramer, E.T., Yao, L., Krasnick, B., et al. (2021). Isotope tracing in adult zebrafish reveals alanine cycling between melanoma and liver. Cell Metab 33, 1493–1504.e1495. 10.1016/j.cmet.2021.04.014.

36. 36. Li, J., Van Vranken, J.G., Pontano Vaites, L., Schweppe, D.K., Huttlin, E.L., Etienne, C., Nandhikonda, P., Viner, R., Robitaille, A.M., Thompson, A.H., et al. (2020). TMTpro reagents: a set of isobaric labeling mass tags enables simultaneous proteome-wide measurements across 16 samples. Nat Methods 17, 399–404. 10.1038/s41592-020-0781-4.

37. Huotari, J., and Helenius, A. (2011). Endosome maturation. EMBO J 30, 3481–3500. 10.1038/emboj.2011.286.

38. Kim, D., Langmead, B., and Salzberg, S.L. (2015). HISAT: a fast spliced aligner with low memory requirements. Nat Methods 12, 357–360. 10.1038/nmeth.3317.

39. Liao, Y., Smyth, G.K., and Shi, W. (2014). featureCounts: an efficient general purpose program for assigning sequence reads to genomic features. Bioinformatics 30, 923–930. 10.1093/bioinformatics/btt656.

40. Love, M.I., Huber, W., and Anders, S. (2014). Moderated estimation of fold change and dispersion for RNA-seq data with DESeq2. Genome Biol 15, 550. 10.1186/s13059-014-0550-8.

41. Ferguson, J.L., and Shive, H.R. (2019). Sequential Immunofluorescence and Immunohistochemistry on Cryosectioned Zebrafish Embryos. J Vis Exp. 10.3791/59344.

42. He, C., and Klionsky, D.J. (2010). Analyzing autophagy in zebrafish. Autophagy 6, 642–644. 10.4161/auto.6.5.12092.

