## Supplementary data for "Chaperone AIP Couples mTORC1 Activation and Catabolic Metabolism During Neonatal Development"

^1^Centre for Endocrinology, Faculty of Medicine and Dentistry, Queen Mary University of London ^2^School of Biological and Behavioural Sciences, Queen Mary University of London, UK. ^3^The Francis Crick Institute, London ^4^Translational Medicine and Therapeutics, ^5^Barts Cancer Institute, Faculty of Medicine and Dentistry, Queen Mary University of London, ^6^Department of Paediatrics, Children's Medical Centre, Landspitali University Hospital, Reykjavik, Iceland, ^7^Department of Neonatology, University Hospital, Leuven, Belgium, ^8^Center for Human Genetics University Hospitals Leuven and Laboratory for the Genetics of Cognition, University of Leuven, Leuven, Belgium, ^9^Research Centre Medical Genetics, Moscow, Russia, ^10^Great Ormond Street Hospital for Children, University College London, London, ^11^Department of Cell and Developmental Biology, University College London, ^12^Biochemical Pharmacology, Faculty of Medicine and Dentistry, Queen Mary University of London, ^13^Faculty of Medicine, University of Iceland, Reykjavik, Iceland ^14^Department of Genetics and Molecular Medicine, Landspitali University Hospital, Reykjavík, Iceland, ^15^Department of Genetic Medicine, Johns Hopkins University, Baltimore, MD, USA, ^16^School of Life Sciences, University of Westminster, London, UK

† These two authors contributed equally

^§^ Current address: Department of Medical and Molecular Genetics, Faculty of Life Sciences and Medicine, King’s College London, London, United Kingdom

**Supplementary material**

#### Patients

Clinical parameters are detailed in **Supplementary Table S1.**

Four of the five patients required neonatal intensive unit care after birth due to extreme weight loss, with failure to thrive despite full nutritional support. All had persistent tachycardia, profuse diarrhea and mild hyponatremia, with pronounced metabolic acidosis. They had intermittent hypoglycaemia (lowest value in patient FCM1 2.86mmol/l), as well as intermittent hypertonia, and gastrointestinal bleeding with anemia in three patients. Two patients had septal atrial defects and one persistent ductus arteriosus and systolic murmur. It should be noted that *Aip* deficient mice have heart developmental abnormalities including septal defects and patent ductus arteriosus^1^. Two patients had some dysmorphic features: FBM1 was noted to have a long and slender body, long fingers and toes, while FCM1 had a triangular face with dysplastic low-set ears, funnel chest, and later craniosynostosis with a small head circumference. All patients had a very low BMI (**Table S1**).

During hospitalization, the main care challenges included feeding problems with recurrent sub-obstruction with subsequent ileostomy formation in one child, diarrhea, failure to thrive despite total parenteral nutrition (TPN) support, episodes of elevated blood pressure (up to 130/80mm Hg), tachycardia (up to 160/minute). One of the major challenges for all five patients was to deal with episodes of unexplained hyperthermia: axillary temperatures ranged usually between 36.5°C and 39.8°C with no sign of infection (**Fig S1A**). This was repeatedly investigated by microbiological assessment of various body fluids, but without positive results, and treated empirically with antibiotics; however, the episodes were not accompanied with elevated leukocytes or CRP, and screenings including bacterial and viral serology were also negative. Occasionally, a higher temperature correlated with an elevated CRP, suggesting true short-term infection, but this was an uncommon finding. All patients showed general high levels of irritability and agitation, with episodes of monotonous loud crying, hyperesthesia and severe hyperhidrosis. They demonstrated arching with head tilted back, seemingly in pain and requiring intermittent sedation. Circulating catecholamines were normal in all 5 patients. Anemia was present in all patients: FCM1 received erythropoietin treatment. Micropolyadenia was noted in two of the 5 children. Hepatomegaly was seen in three patients (1.5cm in FCM1). Cardiac ultrasound identified left ventricular hypertrophy in two children.

*Biochemistry*

Low thyroid stimulating hormone (TSH) levels with a normal free thyroxine (fT4) levels were noted in all children, with raised aldosterone, DHEAS, testosterone and prolactin levels in the one where this was tested. Serum calcium levels were high normal or high (highest calcium levels were: total calcium 3.43mmol/l [2.25-2.75] in FCM1). Hypercalciuria was present, phosphate level was high or normal (range in FCM1 was 1.98-2.56mmol/l [1.45-1.78]) and parathormone was normal or low. In some patients, vitamin D level was supplemented initially as routine but stopped when hypercalcemia was noted. Vitamin D and 1,25 Vitamin D levels were normal. Abdominal ultrasound identified nephrocalcinosis in all the children from the age of 6 weeks, and one of them had duplication of the collecting system on one of the kidneys.

*Imaging*

MRI of the brain revealed no structural abnormalities although there were potential signs of early demyelination in one case. EEG and EMG studies were both within normal limit. FBM1 and FCM1 had bilateral hydrocele. FCM1 had inguinal hernia and underwent surgery at the age of 3 months.

*Therapy*

Four patients received treatment with partial/total parenteral nutrition, repeated courses of antibiotics, erythropoietin, ACE inhibitors, calcium channel blockers, beta-adrenoceptor blockers. Prednisolone was attempted to reduce calcium with biochemical success (high normal calcium and PTH rise achieved), but no clinical improvement. Family A patients had different therapies aiming at increasing caloric intake (parenteral nutrition, insulin therapy with high glucose load, pancreatic enzyme therapy) and decreasing the hypermetabolic state (sedatives, beta blockers, analgesics) without clinical response.

*Outcomes*

FAM1 died age 8 month weighing 4285g. FAM2 passed away at age 10 months old, at that time, he weighed 4090g, with a length of 62.0cm. FBM1 is alive on TPN, has normal IGF-1 and prolactin levels, body height is between 90-97 centile with body weight 25-50 centile. He has moderate intellectual disability with relatively severe speech delay and a diagnosis of autism spectrum disorder. Body temperature elevations ceased around age 11 months. FCM1 had his last admission age 10 months, when he had height at 47 percentile (71cm), weight <0.1 percentile (6.15kg), head circumference <0.1 percentile (41cm), he was lost to follow-up after discharge at 10.5 months of age. FDM1 at the age 14 months has increased muscle tone in the limbs, high popliteal reflexes, and a Babinski sign on the left side. Mental development is delayed but he can pronounce a few words at age 20 months, and his physical condition is stable with slow weight gain, body weight -2.26 standard deviation. His body temperature has remained normal after 14 months of age.

#### Table S1 Clinical parameters of the patients

|  | **Family A**  **FAM1** | **Family A**  **FAM2** | **Family B**  **FBM1** | **Family C**  **FCM1** | **Family D**  **FDM1** |
| --- | --- | --- | --- | --- | --- |
| **Sex** | Female; XX | Male; XY | Male; XY | Male; XY | Male; XY |
| **Ethnicity** | Turkish | Turkish | Icelandic | Ossetian | Ossetian |
| **Mutation in *AIP* gene** | NM_003977.2: c.62G>A | NM_003977.2: c.62G>A | NM_003977.2: c.827C>A | NM_003977.2: c.347_373 deletion | NM_003977.2: c.346_372 deletion |
| **Expected protein change** | NP_003968.2: p.Gly21Asp | NP_003968.2: p.Gly21Asp | NP_003968.2: p.Ala276Glu | NP_003968.2: p.Glu116_Val124del | NP_003968.2: p.Glu116_Val124del |
| **Current status** | Died 8 months | Died 10 months | Alive, closely followed | Lost for follow-up at 10.5 months | Alive, closely followed |
| **Birth weight, length** | 1695 grams, 44.2 cm | 2430 grams, 48.5 cm | 2426 grams, 50 cm | 3100 grams, 49 cm | 3350 grams, 52 cm |
| **Body weight** | 4285 grams at 8 months | 4090 grams at 10 months | 8820 grams at 8 months | 6710 grams at 10 months | 7800 grams at 14 months |
| **Dysmorphic features** | None | None | Long and slender body and fingers and toes, some loose skin of face, funnel chest deformity | Triangular facial shape  Funnel chest deformity  Early closure of cranial vault sutures | None |
| **Pregnancy history** | Born at 31 weeks  Premature rupture of membranes | Fetal tachycardia Polyhydramnios  Born at 36 weeks after induction | Mother had pre-eclampsia requiring early delivery at 35 weeks 4 days. From 30 weeks high blood pressure treated with labetalol and from week 35 preeclampsia, treated with magnesium for 24 hours prior to delivery | Week 37.5 abruption of placenta  Caesarean section | No complications, born at 38 weeks |
| **Genetic testing** | Standard karyotype, whole exome sequencing | Whole exome sequencing | Microarray normal, spinal muscular atrophy testing normal, panel for vitamin D metabolism genes normal, whole genome sequencing | Whole exome sequencing, CYP24A1 heterozygote VUS identified | Whole exome sequencing |
| **Cardiovascular** | Tachycardia, hypertension, large atrial septal defect, L>R shunt, beta blocker for tachycardia and hypertension | Tachycardia,  hypertension beta blocker for tachycardia and hypertension | Tachycardia, hypertension, mild hypertrophy on echocardiography soon after birth, ongoing. On enalapril and propranolol for hypertension and tachycardia | Tachycardia, hypertension, atrial septal defect (hemodynamically insignificant), open arterial duct, left ventricular hypertrophy.  On ACE inhibitor, calcium channel blocker, beta blocker for tachycardia and hypertension | Tachycardia, normal echocardiography, normal blood pressure |
| **Pulmonary** | Normal ventilation, tachypnoea during episodes of hyperthermia | Normal ventilation, tachypnoea during episodes of hyperthermia | Normal ventilation, tachypnoea during episodes of hyperthermia | Apnea, bronchitis | Repeated bronchitis |
| **Gastrointestinal** | Episodes of diarrhea, fat in stool normal, biopsy: non-representative sample, obstruction, high calorie total parenteral nutrition (TPN), needed ileostomy age 18 days | Gastrointestinal bleeding age 7 days, episodes of diarrhea, obstruction, high calorie TPN nutrition | Chronic secretory diarrhea, on long-term TPN nutrition, gastroscopy and colonoscopy normal, gut biopsies no specific abnormality found | Episodes of dehydration, diarrhea | Hepatomegaly, splenomegaly, episodes of poorly digested stool (1-2/day) |
| **Renal** | Polyuria, aminoaciduria, hypercalciuria, normocalcemia | Polyuria, medullar nephrocalcinosis with hypercalciuria, normocalcemia | Hypercalcemia and hypercalciuria, nephrocalcinosis, kidney stones. Treated with low calcium TPN formula (70% of normal).  Bisphosphonate trial lowered blood calcium on 2 occasions of broken bones with minimal/no trauma | Enlarged kidney, hypercalcemia (max value 3.43mmol/l), hypercalciuria, nephrocalcinosis | Hyperechoic inclusions in kidneys, mild hypercalcemia (0.79mmol/l) |
| **Kidney ultrasound** | Increased reflectivity kidney parenchyma, enlarged adrenal glands | Nephrocalcinosis | Nephrocalcinosis | Enlarged kidney Hyperechogenic kidneys  Abnormal vas deferens morphology | Hyperechogenic kidneys |
| **Neurology** | Episodes of increased tone, irritability and discomfort | Episodes of increased tone, irritability and discomfort | Episodes of increased tone, irritability and discomfort.  Has developmental delay (speech and gross motor), formally diagnosed with autism.  Large motor skills behind | Dystonia, encephalopathy, pronounced anxiety (with head tilting), delayed ability to sit, gross motor development | Episodes of increased muscle tone, rapid reflexes, positive Babinski on one side, motor developmental delay, cannot stand or sit at 14 months of age |
| **Temperature** | Episodes of hyperthermia >40°C | Episodes of hyperthermia >40°C | Episodes of hyperthermia, less noticeable after 11 months of age | Episodes of hyperthermia, 38-39°C | Episodes (~monthly) of hyperthermia (38-39°C) until 14 months of age. Subsequently, these episodes became less frequent |
| **Brain MRI** | Delayed myelination of the crus posteriors of the capsula interna | Repeatedly mild ventriculomegaly | Normal | Normal | Not performed |
| **Liver** | Prominent liver, normal intensity | Normal | Normal | Hepatomegaly, elevated hepatic transaminase | Hepatomegaly, transiently elevated liver enzymes |
| **Infection** | Negative | Negative | Low immunoglobulins, on iv immunoglobulins from age 1.5 | Bacteriuria | Repeated bronchitis |
| **Hematology** | Iron-deficiency anemia | Anemia | Anemia, bone marrow biopsy normal, iron infusions needed 3x until age 4.5y | Anemia | Anemia, thrombocytosis |
| **Endocrinology** | Episodes of hyperhidrosis, normal catecholamine, VIP and chromogranin secretion,  low TSH, mild increased aldosterone,  normal DHEAS and testosterone  normal PTH  normal GH and IGF-1,  increased prolactin, intermittent hypoglycaemia | Episodes of hyperhidrosis,  normal catecholamine and cortisol secretion, low TSH, low  1,25-hydroxy VitD, normal PTH,  high-normal GH and IGF-1, intermittent hypoglycaemia | Episodes of hyperhidrosis, on cooling mattress and fan, body covered with  sweat droplets on skin.  Normal urine metanephrins.  Normal thyroid function, normal prolactin, FSH, LH and cortisol.  normal insulin, low PTH, normal 1,25-hydroxy VitD.  Despite normal GH and IGF-1, 2SD above mean for height since age 2 years.  Intermittent hypoglycaemia | Episodes of hyperhidrosis,  normal catecholamines, low TSH,  low PTH,  height 47 percentile while weight <0.1 percentile at age 10m, episodes of hypoglycaemia | Episodes of hyperhidrosis,  low TSH, normal freeT4,  normal 25-OH VitD, GH and IGF-1,  episodes of hypoglycaemia |
| **Immune** | Normal | Normal | Lymphadenopathy (axilla, neck) on both sides, biopsy not concerning | Micropolyadenia (6-9 groups of the increased lymph nodes), decreased antibody level in blood | Micropolyadenia, elevated IL8 levels |
| **Metabolic** | Electrolyte disturbance because of diarrhea; amino acids plasma and urine: normal; organic acids urine- normal; long chain fatty acids: normal; low free carnitine (11 µmol/l [19 – 69]) | Electrolyte disturbance because of diarrhea; normal amino acids in plasma, CSF and urine, normal urine organic acids,  normal sialotransferrins, normal long chain fatty acids,  normal mitochondrial enzymes | Normal plasma amino acids, lactate, acylcarnitine, ammonium highest at 83μM/l | Electrolyte disturbance because of diarrhea,  normal fatty acids,  normal amino acids | Normal amino acid and fatty acid levels,  normal lactate and ammonium |
| **Treatments received** | TPN, steroids, tacrolimus, sedatives, analgesics, beta blockers | TPN, steroids, tacrolimus, sedatives, analgesics, beta blockers | TPN, beta blockers, ACE inhibitors | TPN, repeated antibiotics, oral nutritional support, erythropoietin, ACE inhibitor, calcium channel blocker, beta blocker | Repeated antibiotics, oral nutritional supplements |
| **Comments** |  |  | Gets tired quickly, then lies on the floor | The family is lost from follow-up | Same ethnic background and geographical area as Family C |

**Table S2 Allele frequency of the three identified AIP variants in various databases**

#### (these data do not include the five probands and their parents)

|  | **c.62G>A; p.Gly21Asp**  11-67483220-G-A  (GRCh38) | **c.827C>A; p.Ala276Glu**  11-67490827-C-A  (GRCh38) | **c.346_372del; p.Glu116_Val124del** |
| --- | --- | --- | --- |
| **Alleles tested in gnomAD** | 1614206 | - | - |
| **Alleles found in gnomAD** | 5 heterozygote (2 non-Finnish European, 3 far Eastern)  0 homozygote | Not identified | Not identified |
| **Allele frequency in gnomAd** | 0.000003097 | - | - |
| **Alleles tested in Iceland** |  | 319218 | - |
| **Alleles found in Iceland** | Not identified | 27 heterozygote  0 homozygote | Not identified |
| **Allele frequency in Iceland tested population** |  | 0.00008458 |  |
| **Alleles tested in Russia Genetic project** |  |  | 78985 |
| **Alleles tested in Russia Genetic project** | Not identified | Not identified | 3 heterozygote (all 3 from the same geographical region as probands)  0 homozygote |
| **Allele frequency in Russia tested population** |  |  | 0.00003798 |

### Supplementary Figure legends

**Supplementary Figure 1**

Axillary temperature profile over a period of 4 months from patient FBM1 showing spikes in temperature coinciding with periodic fevers (A). Location of the variants in AIP (B). Family A and Family B AIP variants half-life as determined by cycloheximide (CHX) treatment of HEK cells transfected with plasmids containing *AIP* variants and analyzed by Western blotting. Expression of AIP was normalized to a loading control and to expression at time 0. Graphs show the mean ± SEM from at least two independent experiments (C).

**Supplementary Figure 2**

WT and *Aip* KO MEFs were starved overnight in DMEM containing 0.1% serum and stimulated with 5ng/ml epidermal growth factor (EGF) (A), 10% fetal bovine serum (FBS) (B) and 100μM sphingosine-1 phosphate (S1P) (C) for the indicated time points. p-AKT, p-ERK and p-S6K1 expression was examined by Western blotting. Graphs show the mean ± SEM from at least three independent experiments.

**Supplementary Figure 3**

Viability of WT and *Aip* KO MEFs cultured in EBSS and EBSS plus bafilomycin (Bafilo) (A). Expression of LC3 and ubiquitin in WT, *Aip* KO MEFs and *Aip* KO “rescue” MEFs (B). Number of PI3P^+^ (C), Wipi2^+^/PI4P^+^ (D), PI4K3β^+^ puncta (E) in WT and *Aip* KO MEFs under starved (2 hours EBSS) conditions. Graphs show the mean ± SEM from at least two independent experiments. Students un-paired, paired *t*-test (A, D, E) and one-way ANOVA with Tukey’s multiple comparison test was used for analysis (B) and 2-way ANOVA with Tukey’s multiple comparison test was used for analysis (C, E).

**Supplementary Figure 4**

TFEB expression determined by Western blotting under fed and starved (2 hours EBSS) conditions in WT and *Aip* KO MEFs (A). Expression of TFE3 in fed and starved WT and *Aip*-KO-MEFs, A.U. arbitrary units (B). Autophagy gene expression in WT and *Aip* KO MEFs under fed conditions determined by qPCR (C). Graphs show the mean ± SEM from at least two independent experiments. Students un-paired *t*-test (A-C) was used for analysis.

**Supplementary Figure 5**

Induction of autophagy by culturing healthy control (HC) and AIPd-PDFs (FAM1, FBM1 and FDM1) in EBSS determined by immunofluorescence for Wipi2 (A) and Western blotting for LC3 and p62 (B). Induction of autophagy by treating healthy control (HC) and AIPd-PDFs with rapamycin (Rapa) 100nM and bafilomycin (Bafilo) 100nM for 2 hours (C). Expression of autophagy genes in HC and AIPd-PDFs (FBM1) under fed conditions determined by qPCR (D). ­Graphs show the mean ± SEM from at least two independent experiments. 2-way ANOVA with Tukey’s multiple comparison test (A) and Students un-paired *t*-test (D) was used for analysis.

**Supplementary Figure 6**

Electron microscopy of WT and *Aip* KO MEFs, highlighting the autophagosomes (green arrows), endosomes (yellow) and lysosomes (purple) (A). Electron microscopy analysis of HC and AIPd-PDFs (FBM1) with the number of endosomes and autophagosomes determined (B). Graphs show the mean ± SEM from at least two independent experiments. Students un-paired *t*-test (A-B) was used for analysis.

**Supplementary Figure 7**

Heat map of RNA-sequencing data showing relative expression of lysosome genes (from GSEA <https://www.gsea-msigdb.org/gsea/msigdb/index.jsp>) under fed conditions in WT and *Aip* KO MEFs (A). Heat map of RNA-sequencing data showing relative expression of genes involved associated with lysosomal storage diseases. List of genes from <https://panelapp.genomicsengland.co.uk/panels/529/> (B). Expression of NPC2 and GAA in WT and *Aip* KO MEFs normalized to total protein (C). Graphs show the mean ± SEM from at least two independent experiments. Students un-paired *t*-test (C) was used for analysis.

**Supplementary Figure 8**

Glucose import into WT and *Aip* KO MEFs was determined using the fluorescent glucose analogue (2-NBDG) (A) and the relative amount of uniformly labelled glucose (U-^13^C_6_ (m+6)) in WT and *Aip* KO MEFs (B). The rate of glycolysis was measured by treating the cells with sodium oxamate and measuring the ratio of NADH/NAD in WT and *Aip* KO MEFs. Insert shows expression of LDHA in WT and *Aip* KO MEFs by Western blotting (C). Metabolomic analysis using uniformly labelled glucose (U-^13^C_6_) in WT and *Aip* KO MEFs measuring total glycolysis products and flux (m+3) from labelled glucose (D). Relative proportion of amino acids derived from the culture supernatant of WT and *Aip* KO MEFs (E). Relative amounts of glycolysis metabolites from the culture supernatant of WT and *Aip* KO MEFs from uniformly labelled (U-^13^C_6_) glucose (F). Viability of WT and *Aip* KO MEFs and healthy control (HC) and AIPd-PDFs (FBM1) cultured in glucose free media over 48 hours (G). Viability of WT and *Aip* KO MEFs cultured in the presence of glycolysis inhibitor 2-deoxy-D-glucose (2-DG) (10mM) and ATP synthase inhibitor oligomycin (Oligo) (1μM) after 48 hours (H). Graphs show the mean ± SEM from at least two independent experiments. Students un-paired (A-F, H) and 2-way ANOVA with Tukey’s multiple comparison test (G) was used for analysis.

**Supplementary Figure 9**

Relative amounts of TCA cycle intermediates derived from uniformly labelled glucose (U-^13^C_6_) (A-C) from WT and *Aip* KO MEFs. Mitochondrial membrane potential was determined in WT and *Aip* KO MEFs using TMRM (D). Mitochondria volume and membrane potential was determined in WT and *Aip* KO MEFs using MitoTracker red/MitoTracker green (E) and MitoTracker (F) dyes by flow cytometry and median fluorescent intensity (MFI) determined. Mitochondrial NAD/NADH was measured in WT and *Aip* KO MEFs (G) The ATP/ADP ratio was determined in WT and *Aip* KO MEFs (H). Results of at least two independent experiments. Graphs show the mean ± SEM from at least two independent experiments. Students un-paired *t*-test was used for analysis (A, B, D-H).

**Supplementary Figure 10**

WT and *Aip* KO MEFs were treated with uniformly labelled carbon glutamine (U-^13^C_5_) and the relative amounts of TCA intermediates and the relative proportion of oxidative (m+4) and reductive (m+3) glutamine metabolism calculated (A). Using uniformly carbon (U-^13^C_5_) and nitrogen (U-^15^N_2_) labelled glutamine, the relative proportion of in glutamine metabolites from WT and *Aip* KO MEFs (B). Serum amino acids from FAM1 and FBM1 were analyzed and compared against the normal range for these amino acids (C).Graphs show the mean ± SEM from at least two independent experiments. Students un-paired *t*-test was used for analysis (A).

**Supplementary Figure 11**

Comparison of human and zebrafish AIP genomic DNA and AIP structure as determined by Alphafold and Swissfold. The zebrafish AIP protein shares 78.8% identity with the human protein (A). Design and generation of a homozygous *aip* mutant zebrafish line. Strategy used to target *aip* in zebrafish using CRISPR/Cas9 (B). The 29 bp frame shift caused by the deletion leads to a premature stop codon in the PPIase domain, thereby disrupting the downstream C-terminal tetratricopeptide repeat (TPR) domain.

**Supplementary Figure 12**

Metabolic analysis of zebrafish. Relative amounts of glycolysis products (A), acetyl-CoA and TCA cycle intermediates and amino acids derived from uniformly labelled glucose (U-^13^C_6_) (B). Glutaminolysis products derived from labelled glutamine derived from uniformly labelled glutamine (U-^13^C_5_) and (U-^15^N_2_) (C). Students unpaired *t*-test was used for analysis (A-C).

**Supplementary Figure 13**

##### **Healthy Neonates:**

**Nutrient Sufficiency:** In healthy cells, the presence of nutrients and growth factors activates mTORC1 at the lysosome, driving **anabolism, cell growth**, and **protein synthesis**. Amino acids are replenished through **proteasome-mediated protein degradation**, maintaining metabolic balance. Growth factor receptor mediated signaling results in acquisition of nutrients into the cell.

**Nutrient Scarcity:** C**atabolic processes**, particularly **autophagy**, are activated. Cells recycle intracellular proteins to generate amino acids, which sustain **mTORC1 activity** and supports ongoing growth in the absence of external nutrients.

##### **AIP-Deficient Neonates:**

**Nutrient Sufficiency:** Despite the availability of external nutrients, **mTORC1 activation is impaired** due to defective **growth factor signaling** and **proteasome dysfunction**. This results in the **accumulation of ubiquitylated proteins**, disrupting cellular homeostasis and hindering growth and reduced acquisition of nutrients into the cell.

**Nutrient Scarcity:** AIP-deficient cells fail to induce **autophagy**, leading to a severe **shortage of intracellular nutrients, energy depletion**, and further impairment of **mTORC1 signaling**. This failure to adapt metabolically is especially damaging during periods of **nutrient scarcity**, contributing to **postnatal lethality** in AIP-deficient infants.

##### **Summary:**

AIP plays a vital role in **nutrient recycling** and **metabolic adaptation**. Its loss disrupts both anabolic and catabolic pathways, preventing cells from maintaining metabolic balance. This collapse becomes critical under nutrient-scarce conditions, particularly during the neonatal period when rapid metabolic adaptation is essential for survival and sustained organismal growth.

**References**

1.	Lin, B.C., Sullivan, R., Lee, Y., Moran, S., Glover, E., and Bradfield, C.A. (2007). Deletion of the aryl hydrocarbon receptor-associated protein 9 leads to cardiac malformation and embryonic lethality. J Biol Chem *282*, 35924-35932. 10.1074/jbc.M705471200.
